# The emergence of the fractal bronchial tree

**DOI:** 10.1101/2025.01.13.632436

**Authors:** Malte Mederacke, Kevin A. Yamauchi, Nikolaos Doumpas, Laura Schaumann, Jonathan Sperl, Thomas Weikert, D. Merrill Dane, Connie C. W. Hsia, Maurice Pradella, Jens Bremerich, Roman Vetter, Dagmar Iber

## Abstract

The fractal design of the bronchial tree, as described by the Hess–Murray law, facilitates energy-efficient lung ventilation, yet its developmental origins remain unclear. Here, we quantify the rearrangement of the embryonic bronchial tree into its fractal architecture, using SkelePlex, a new neural network-based image processing pipeline, and elucidate the biophysical principles governing its formation. We find that the branch shapes are such that pressure drop, shear, axial and hoop stress are equal across all branches. The seemingly random diameter of sister branches reflects the different number of tips they connect to in the asymmetric lung tree. By analysing lungs after pneumonectomy and those affected by chronic obstructive pulmonary disease (COPD), we show the same principles to persist in adulthood and disease, potentially providing quantitative biomarkers for disease progression and therapeutic guidance.

## Introduction

The lung mediates gas exchange between the blood and the air. Adult bronchial trees in mouse and human are composed of 23–26 dichotomous branch generations, incrementally decreasing in size from the trachea [2, 4, 5]. The architecture of the dichotomously branching bronchial and vascular trees has long been related to energy minimisation [4, 6–8], but the mechanism by which the optimal tree architecture develops in time for the first breath at birth has remained elusive.

In conducting trees, there is a critical trade-off between dead volume in wide tubes and the cost of transport due to flow resistance in thin tubes (Fig. 1a,b) [6–8]. In an idealised, symmetric, dichotomous tree with cylindrical branches, with laminar flow according to the Hagen–Poiseuille equation, the energy expenditure is minimal if the sum of the cubed inner diameter of the two daughter branches, *D*_*z*_, is equal to the diameter cubed of the parent branch: 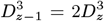. This yields a homothety ratio of the inner branch diameters of subsequent generations *z* (counting from the trachea *z* = 0), given by

**Figure 1:**
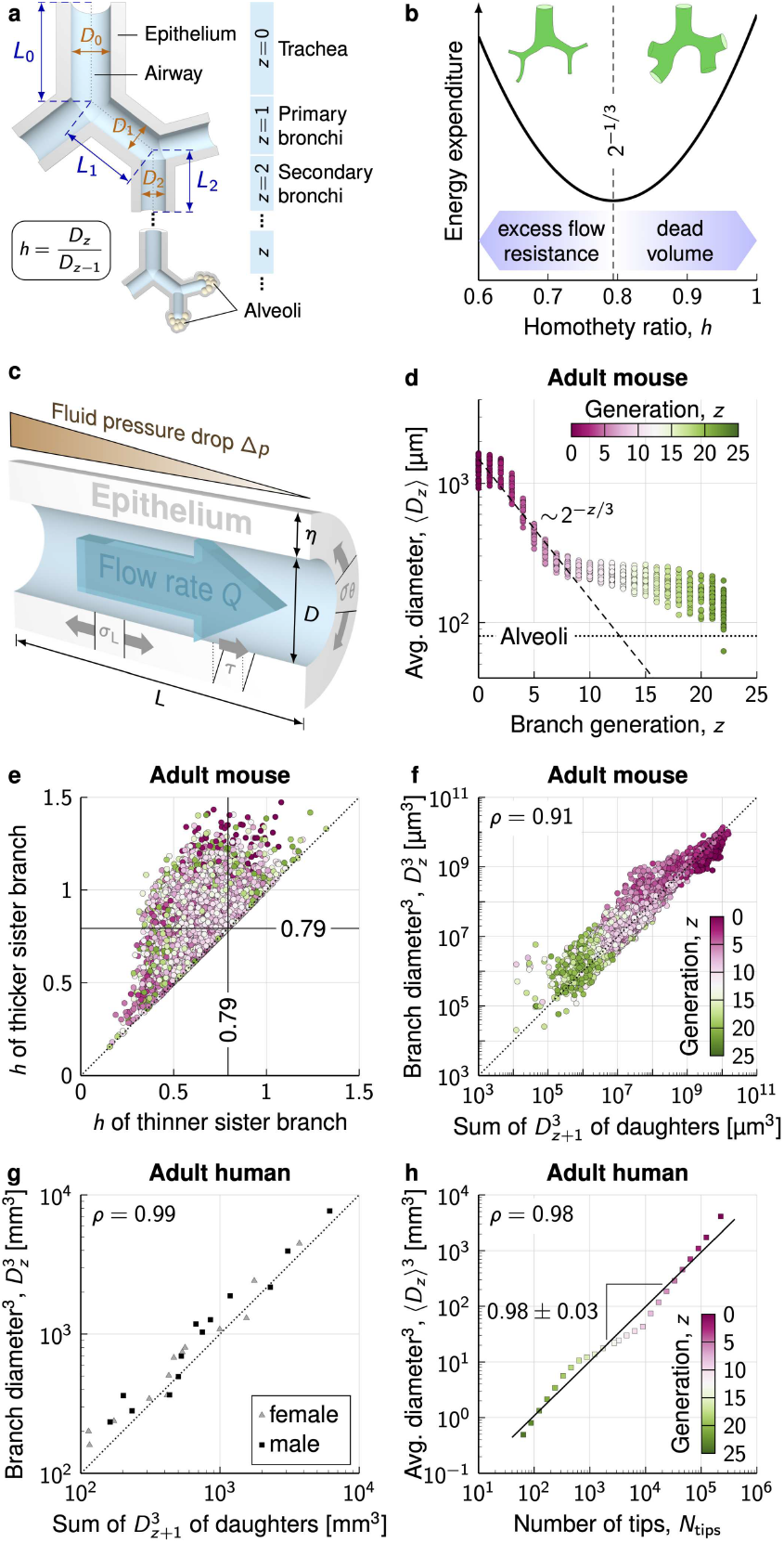
Organisation of the adult bronchial tree. **a**, Overview of the bronchial tree. **b**, Hess–Murray law: A homothety ratio of *h* = 2^−1/3^ permits ventilation with lowest energy expenditure. **c**, Mechanical effects of fluid flow on the epithelial wall (*τ* : shear stress, *σ*_L_: longitudinal stress, *σ*_*θ*_: hoop stress). **d**, The average inner branch diameter per generation largely follows the Hess–Murray law (dashed line) for the first 8 generations *z* in adult mouse lungs. **e**, Homothety ratios *h* of the inner branch diameters of sister branches show that individual branches deviate from the Hess–Murray law. **f**,**g** 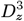 of parent branches vs. the sum of 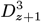 of their daughter branches in the adult mouse lung **(f)**, and the adult female (triangles) and male (squares) human lung. **h**, The average cubed inner branch diameter scales with the number of tips in the adult human lung. *ρ* are Pearson correlation coefficients. Panels **d–f** combine data from *n* = 34 lungs from four different mouse strains [1, 2], panels **g**,**h** show individual human lungs [3]. In **h**, 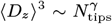 with fitted exponent *γ* = 0.98 ± 0.03 (s.e.).

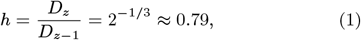

and the branch diameters follow the Hess–Murray law [6, 7]

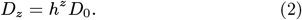

For incompressible fluids, the combined volumetric laminar flow rate must sum up to *Q*_*z*−1_ = 2*Q*_*z*_, resulting in equal wall shear stress, *τ* in all branches (Fig. 1c):

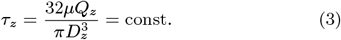

*µ* represents the dynamic viscosity, which is about ten times higher in the lumenal fluid of the developing lung than in water [9]. These relationships are independent of branch shape and angles. Fixed branch shapes, i.e., a constant branch length to diameter ratio, *L*_*z*_/*D*_*z*_, result in an equal pressure drop across all branches [10],

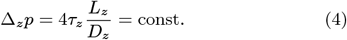

Morphometric measurements of adult human and rodent lungs are impressively close to these theoretical predictions [4, 5, 11–18], as we illustrate by replotting published data [1, 2] from adult mice (Fig. 1d and S1), but there are important deviations. Part of these reflect technical limitations in the quantification, in particular as branches shrink in size [11]. More importantly, however, the lung tree is highly asymmetric [5, 11, 19] such that few daughter branches shrink by the same homothety ratio, and even fewer by *h* = 0.79 (Fig. 1f). In the five human lungs that Weibel analysed sixty years ago, only 8% and 35% of the conjugate branches analysed had equal lengths or diameters, respectively [11]. Thinner sister branches are often longer than their siblings [20, 21]. When considering wider distributions of homothety ratios in each generation, the exponential relationship (Eq. 2, Fig. 1d dashed line) turns into a power law in distal airway generations [22], which fits the various datasets from human, rat, dog, and hamster [4, 11, 13]. The highly asymmetric architecture of the bronchial tree serves an important purpose in that it reduces the average transit time between trachea and alveoli, thereby maximising gas delivery during the brief periods set by pulsatile breathing [23] without increasing flow resistance [24].

Despite the branch asymmetry, the homothety ratios of sister branches are not random. Rather, the sum of the inner diameter cubed of the two daughter branches, 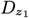 and 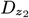, is equal to the diameter cubed of the parent branch (Fig. 1f,g),

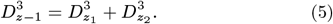

This relationship is termed Murray’s law and follows from the same assumptions that lead to the Hess–Murray law except for branch symmetry. As a consequence of Murray’s law, the branch diameter cubed is proportional to the number of tips (also referred to as leaves in the theoretical literature on tree structures) to which the branch connects [5] (Fig. 1h),

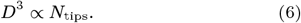

Departures from a cubic relationship, as observed in studies on rabbit lungs, can result from deviations from laminar flow, or reflect limitations in the data [24, 25].

Given the differences between sister branches, the bronchial tree exhibits self-similarity only when considering suitable branch averages (Fig. 1b,d) [24]. This raises the question whether a single mechanism can establish the highly asymmetric, yet highly optimised architecture of the lung on length scales spanning many orders of magnitude. What provides the local guidance cues to cells to generate the globally optimal branch shapes, and ultimately, an optimal tree?

Mouse lung development commences around embryonic day (E)9.5 with the induction of the lung field in the ventral foregut, and proceeds with the emergence of the two main bronchi at E10.5 (Fig. 2) [26]. In the subsequent pseudo-glandular stage that lasts until E16.5, the bronchial tree undergoes multiple rounds of branching morphogenesis [27, 28]. At the canalicular stage, beginning around E17.5, the terminal buds start to narrow [28]. Then, during the saccular stage, from E18.5 to postnatal day (P)5, they form multiple small sacs, which serve as precursors to the alveoli. Alveoli together with respiratory bronchioles form acini, the primary functional unit for gas exchange. As the animal grows, the number of acini remains constant, but their volume expands [29].

**Figure 2:**
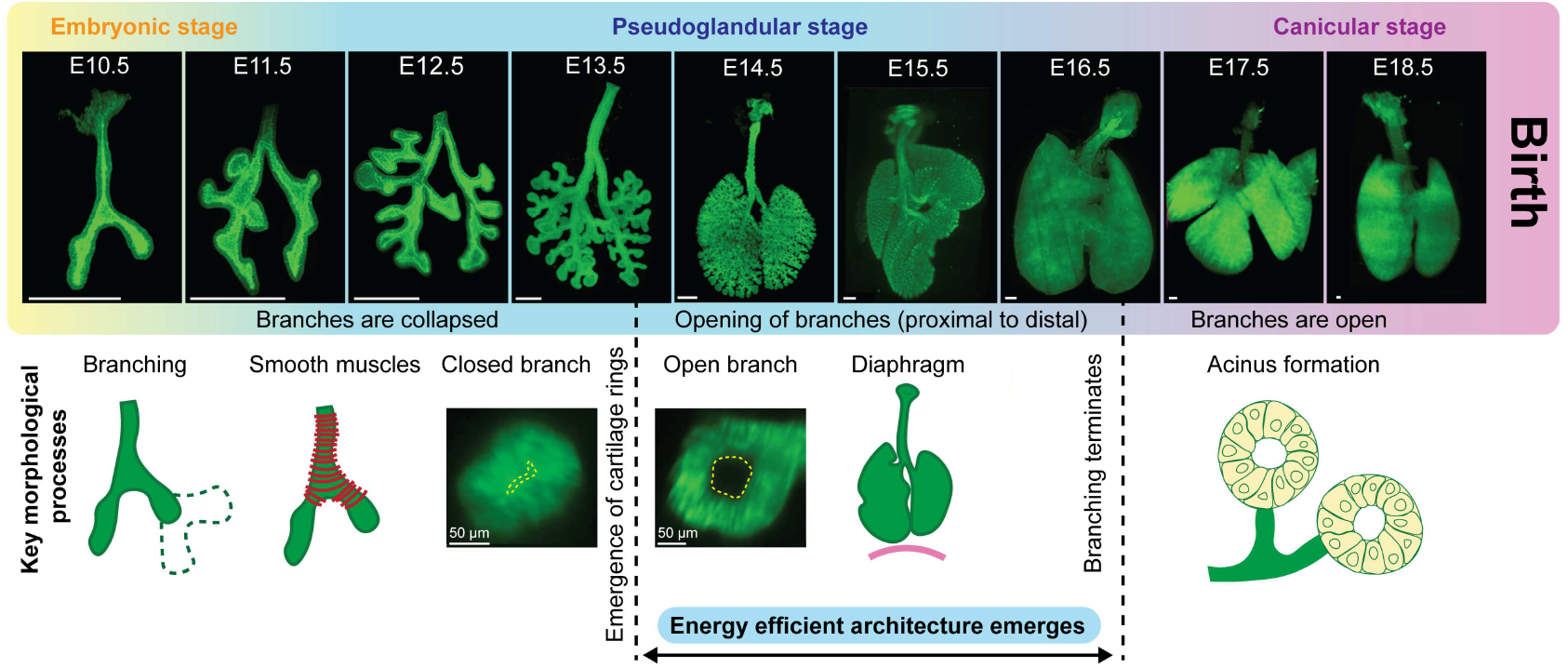
Progression of embryonic lung development. Until E10.5 in embryonic mouse lung development, the lung field develops from the endodermal foregut and the two main bronchi emerge. During the pseudo-glandular phase (E11–16.5), branching morphogenesis defines the number of branches. Once cartilage and smooth muscles stabilise the tubes, the closed branches open up sequentially from proximal to distal. Around E16.5, all branches are open, branching morphogenesis terminates, and the lung transitions into the canicular phase. Here, the first acini form, followed by the development of alveoli postnatally. Scale bars are 500 µm, unless noted differently.

The energy-efficient lung architecture must emerge via extensive branch remodelling because the inner branch diameters remain tiny until emerging smooth muscles and C-shaped cartilage rings support larger fluid-filled lumina from E14.5 onward (Fig. 2, S2) [30, 31]. Fluid-structure interactions have long been suspected to drive the (re-)modelling of biological transportation networks, including blood vessels, plant vasculature, slime mold networks and branched epithelial organs, via branch dilation, constriction, and network pruning [32–36]. Fluid flow is observed in developing lung tubes from the earliest stages of development [30, 37]. At later stages, rhythmic contractions of the diaphragm and chest muscles, called fetal breathing, drives the fluid flow [38, 39]. Lung fluid that leaves through the trachea is either swallowed or enters the amniotic sac [40, 41]. Assuming laminar flow, the inner diameters could, in principle, be adjusted to meet Murray’s law to achieve a common wall shear stress [42, 43]. As E14.5 has remained the latest mouse developmental stage for which a murine bronchial tree has been quantified [44], and only the branch length has been quantified during the pseudo-glandular stages (Carnegie stage (CS)15 and CS23) of developing human lungs [45, 46], it is unknown how and when the inner diameter remodels.

To comprehensively quantify the developing bronchial tree during the key remodelling stages, we developed a computational pipeline, SkelePlex, leveraging advanced image analysis techniques (Fig. 3). SkelePlex was applied to 3D imaging data obtained through state-of-the-art tissue-clearing methods and light-sheet microscopy. Our findings suggest that fluid-structure interactions may provide the local cues to guide the cellular re-arrangements that lead to a globally optimal organ architecture. Specifically, we propose that inner diameter ratios evolve as epithelial tubes expand to maintain a uniform wall shear stress, while wall thickness adjusts to achieve a common hoop stress. By analysing canine lungs after pneumonectomy as well as a large clinical human dataset, we show that these biophysical principles persist into adulthood and help explain structural changes in patients suffering from chronic obstructive pulmonary disease (COPD), potentially offering novel approaches for disease detection and monitoring.

**Figure 3:**
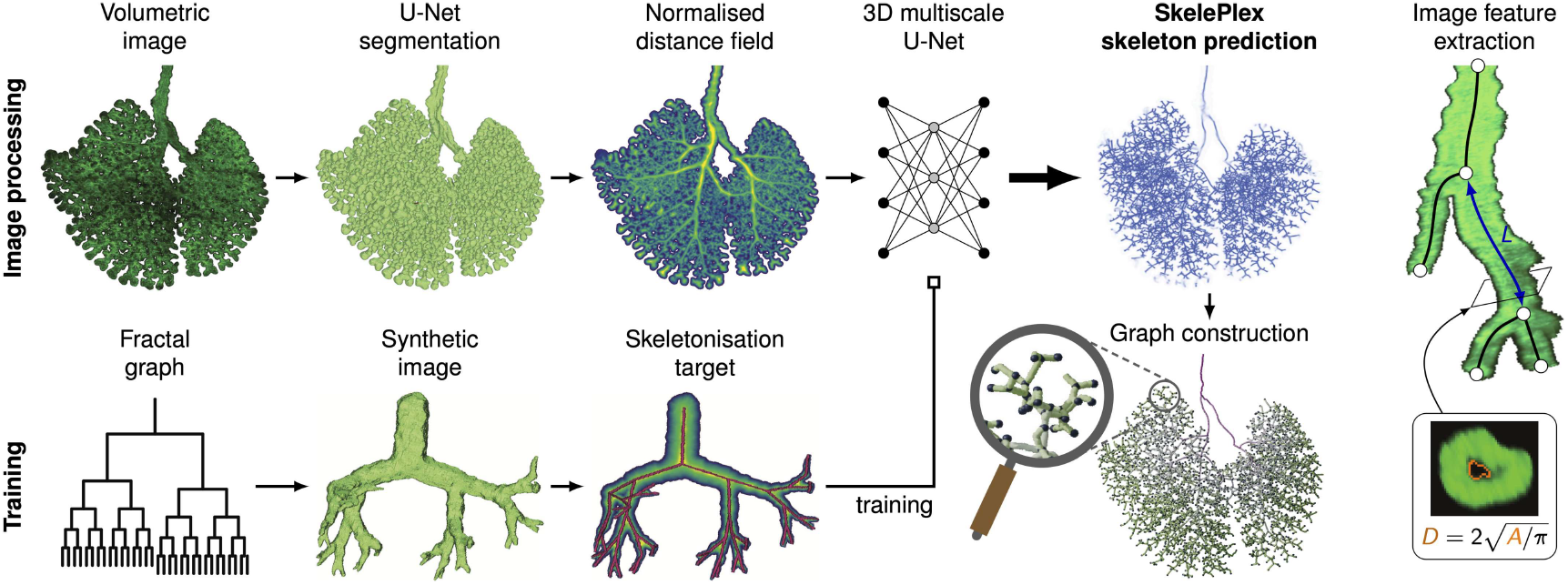
SkelePlex for quantification of bronchial trees. Trained on synthetically generated fractal trees, the deep-learning-based image analysis pipeline SkelePlex skeletonises light-sheet fluorescence microscopy images of whole lungs and measures their morphology and bronchial tree architecture.

## Results

### SkelePlex quantifies lung architecture across scales and modalities

We exploited a transgenic mouse line that expresses membrane-tagged green fluorescent protein (GFP) in the lung epithelium to obtain a 3D time series of murine embryonic lungs at daily intervals (Fig. 2). To analyse and quantify the complex branching structure of the bronchial tree from these 3D images, we developed an open-source end-to-end pipeline named SkelePlex (Fig. 3). SkelePlex integrates advanced preprocessing and synthetic training data to address three key challenges: (1) accommodating diverse imaging modalities across organisms and conditions, (2) overcoming limited ground truth data for model training, and (3) handling the lung’s multiscale architecture, spanning multiple orders of magnitude in length scale.

In the first stage of the pipeline, the branches are segmented based on the fluorescent membrane signal using a 3D U-Net. Then, a multiscale 3D U-Net is used to predict the skeleton from the normalized Euclidean distance transform of the segmentation. By separating the segmentation and skeletonisation steps, the skeletonisation network can be trained with synthetic data, which overcomes the need for laboriously hand-curated lung skeletons to generate training data. The same skeletonisation network can then be used across across different imaging modalities. To handle the lung’s varying scales, SkelePlex preprocesses images using distance transformation and local normalisation, ensuring consistent branch intensity patterns across scales. SkelePlex then extracts a graph representation of the branching architecture whose nodes are the branch points and whose edges are splines fitted to the branch skeleton. Using this graph representation, SkelePlex measures quantitative features such as the branch length, diameter, wall thickness, and angle. We have successfully quantified lung architectures in this work from fluorescence microscopy and computer tomography (CT) imaging.

### Emergence of the self-similar organisation in the embryonic mouse lung

Using SkelePlex, we measured the morphological features of embryonic mouse lungs throughout the pseudo-glandular stage up to E16.5, covering the full extent of branching morphogenesis (Fig. 4a,b). Consistent with the importance of the asymmetric lung tree architecture [23, 24], the fraction of branches per side and per lobe remains relatively constant from the initial emergence of the two main bronchi at E10.5, through the subdivision of the right lung into four lobes at E12.5, up to the completion of branching morphogenesis at E16.5 (Fig. 4c). The degree of asymmetry thus develops similarly in the entire lung.

**Figure 4:**
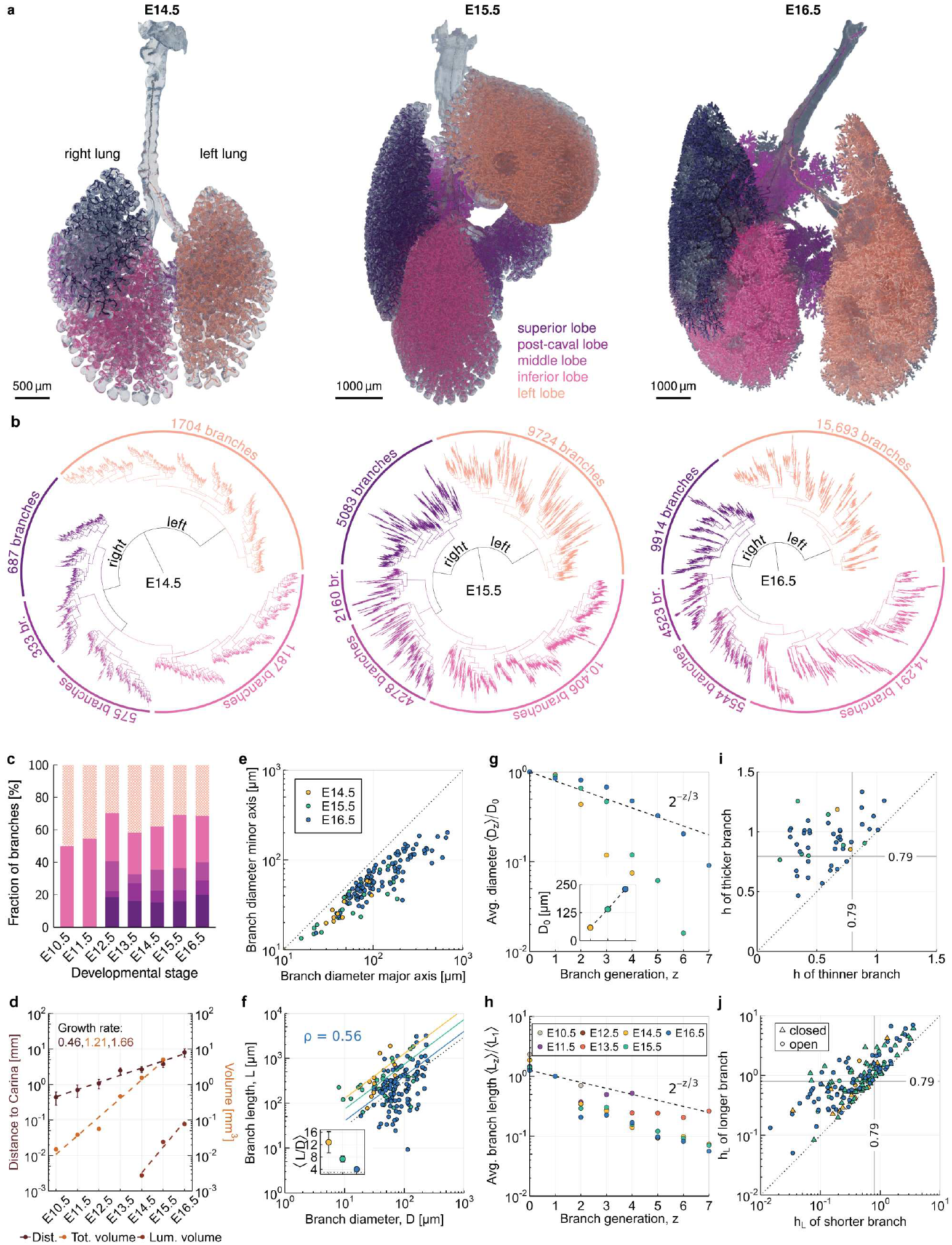
Emergence of the self-similar organisation in the embryonic mouse lung. **a**, Surface reconstructions of light-sheet-imaged embryonic mouse lung epithelia (transparent) whose lobes (coloured) were skeletonised with SkelePlex. **b**, Corresponding hierarchic organisation of the branches in the dichotomous bronchial trees. **c**, Fraction of branches per lobe over developmental time. **d**, Average distance from the tips to the carina, total luminal volume and total epithelial volume over developmental time. Growth rates are in d^−1^. **e**, Length of the longest axis and its orthogonal axis as a measure of circularity of the branch lumen. **f**, Branch length-to-diameter ratio. Solid lines are linear fits. *ρ* is the Pearson correlation coefficient at E16.5. Inset: The geometric mean of *L*_*z*_/*D*_*z*_ from E14.5–16.5 approaches 3 (dotted line). Error bars are s.e. **g**,**h**, Inner branch diameters **(g)** and branch lengths **(h)** against the branch generations, starting at the trachea (*z* = 0). Dashed lines represent the Hess–Murray law. **i**,**j**, Homothety ratios *h* = *D*_*z*_/*D*_*z*+1_ and *h*_*L*_ = *L*_*z*_*/L*_*z*+1_ of the inner branch diameters (**i**) and lengths (**j**) of sister branches.

The total epithelial volume and the average distance of the tips to the carina increase exponentially over developmental time (Fig. 4d). The branch lumen remains narrow (*D*_*z*_ ≈ 0) until E13.5 [30] (Fig. S2), but once the branches open up, also the total branch lumen increases exponentially over time (Fig. 4d). The branches open sequentially in a proximal-to-distal direction. By E14.5, open luminal spaces are observable and quantifiable in at least some branches of the first five generations, but only the first pair of sister branches has developed a visible lumen. By E15.5, open branches extend to the first seven generations, and by E16.5, all branch generations exhibit open lumens. The branches are largely circular, though a more elliptic shape emerges the wider the lumina become (Fig. 4e). By E16.5, the average length-to-diameter ratio approaches 3 (Fig. 4f), the average homothety ratios of the lumenal diameters approach *h* = 2^−1/3^ over the first seven generations (Fig. 4g, Fig. S3a), while the branch lengths deviate from *h*_*L*_ = *L*_*z*_*/L*_*z*−1_ =2^−1/3^(Fig. 4h, Fig. S3b). We conclude that the energy-efficient design emerges after branches have formed, but while the lung is still growing.

### Uniform shear stress in the bronchial tree

Much as in the adult lung, individual branches of the embryonic lung deviate substantially from the average, such that barely any individual branch follows the homothety ratio of the Hess–Murray law, or is three times longer than wide, and few sister branches have equal length and diameter (Fig. 4f,i,j, Fig. S3b–d). Despite the enormous variability in the branch diameters (Fig. 4i, Fig. S3a), we find the cubed diameter of the parent branch to be equal to the sum of the cubed diameters of its daughter branches (Fig. 5a). As the lung liquid is incompressible and the volume flow rate of a parent branch must thus be equal to the sum of its two daughter branches, this implies equal wall shear stress *τ* in all branches (Eq. 3). The variations in tube diameter along each branch are small, and thus do not substantially increase the net shear stress over that expected for the mean diameter (Fig. S4). Consistent with Murray’s law, we further find *D*^3^ ∝ *N*_tips_ in the mouse embryonic lung by E16.5 (Fig. 5b), the developmental time point by which all branches have opened up and fluid can flow freely.

**Figure 5:**
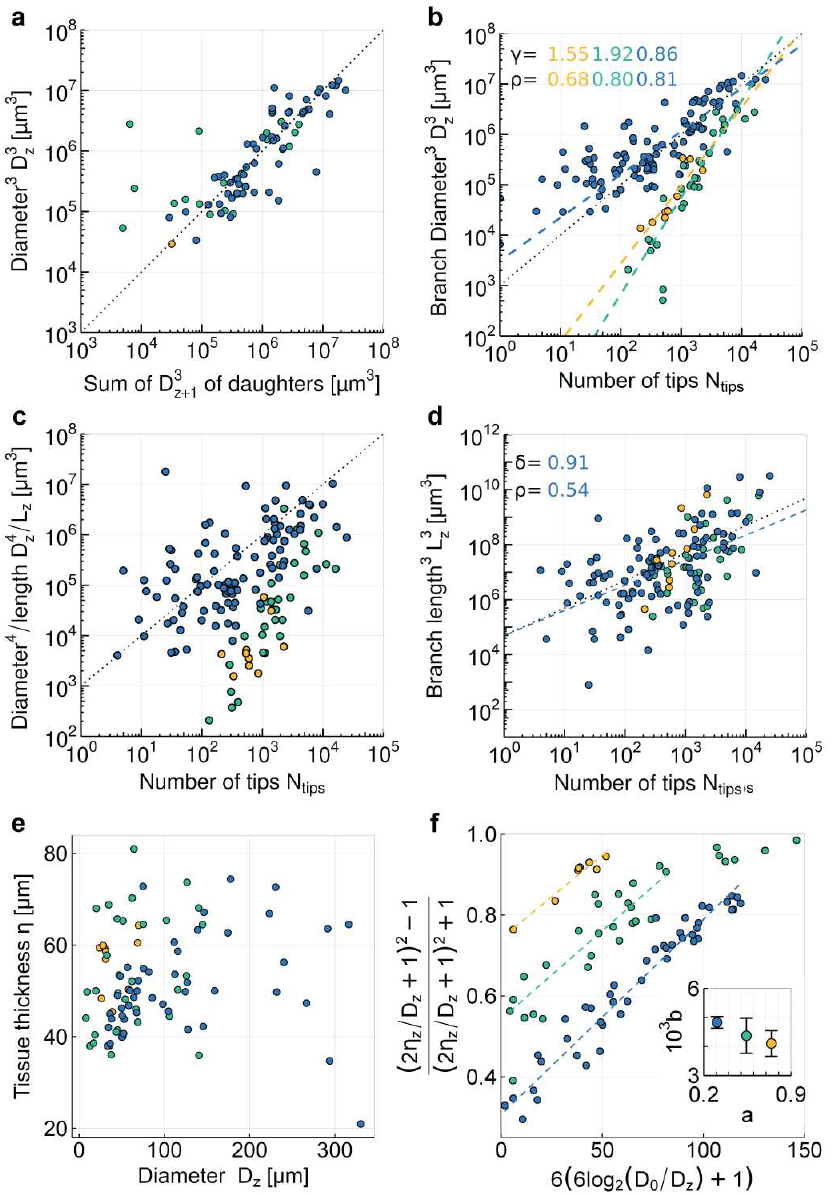
Emergence of the self-similar, but variable lung architecture as a result of fluid-structure interactions. **a**, 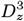of parent branches vs. the sum of 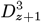 of their daughter branches in the embryonic mouse lung at E14.5 (yellow), E15.5 (green) and E16.5 (blue). **b**, The cubed branch diameter scales with the number of tips in the E16.5 mouse lung. The exponent *γ* in 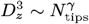 approaches 1 from E14.5 to E16.5. 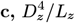 scales with the number of tips in the E16.5 mouse lung. **d**, The cubed branch length scales with the number of tips in the E16.5 mouse lung. The exponent *δ* in 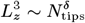 approaches 1 (dotted line) at E16.5 (blue). **e**,**f**, Relationship between airway diameter and epithelial wall thickness in developing mouse lungs. Fits with Eq. S15. Inset: The inferred biophysical parameters *a, b*, as listed in Table S1. Error bars are s.e.

### Constant pressure drop across each branch

As the branch diameters increasingly follow Murray’s law (Eq. 5), we observe that the branch lengths and diameters become increasingly correlated, i.e., *L_z_* ∝ *D_z_* (Fig. 4f), and 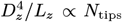 (Fig. 5c). The pressure drop across branches (Eq. 4) thus becomes increasingly similar. This relationship likely emerges because of space limitations. As the branch lengthens, there is more space for additional branching and tip formation, resulting in 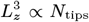 (Fig. 5d). Enlargement of the chest cavity is met by expansion through branching morphogenesis and branch lengthening, again leading to the observed proportions.

### Uniform hoop stress in the bronchial tree

The pressure gradient imposes a stress gradient on the epithelial tube wall (Fig. 1c). As the epithelial wall thickness, *η*, is of the same order as the branch diameter, *D* (Fig. 5e, Fig. S5), we can apply the thick-walled cylinder approximation (Supplementary Text) to analyse the stress distribution within the epithelial wall caused by the pressure difference between the interior and exterior of the epithelial tube (*p* and *p*^out^, respectively). The hoop stress in circumferential direction is given as

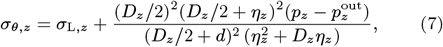

where the axial stress,

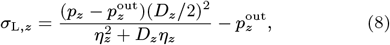

describes the stress in longitudinal direction in the tube wall.

Vertical intrapleural pressure gradients have been reported for adult lungs [47–49], and could cause 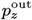 to change with vertical distance from the tracheal opening (Supplementary Text). However, determining vertical distances in embryonic lungs is challenging because the lobes can shift during lung extraction. Therefore, for embryonic lungs, we assume 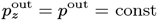. Adult lungs are ventilated by creating a negative pressure within the chest cavity, which draws air in passively. During exhalation, the chest cavity decreases in volume, creating positive pressure that pushes air out of the lungs. Similarly, fluid flow is bidirectional in the developing lung, though there is a net outflow of lung liquid that increases over developmental time [40, 41, 50, 51]. While airflow is likely turbulent over the first branch generations of the human lung, fluid flow in the developing lung is likely laminar throughout, given its higher viscosity and smaller tubular diameter [52]. Assuming laminar flow, there is either a linearly increasing or decreasing pressure gradient from the tracheal opening. The average pressure in generation *z* can then be approximated as

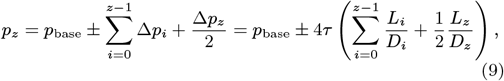

where *p*_base_ is the pressure at the outlet of the trachea, the airway opening pressure. The positive or negative sign reflects the direction of the pressure gradient. Using *L*_*i*_/*D*_*i*_ ≈ ⟨*L*/*D*⟩ ≈ 3 and *z* = 3 log_2_(*D*_0_/*D*_*z*_), this simplifies to

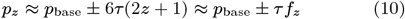

where

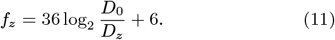

Eqs. 10,11 require only local branch information and can be used also when the tree reconstruction is missing. In combination, Eqs. 7–11 yield relationships between wall thickness and branch diameter at the luminal surface (Supplementary Text),

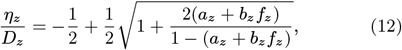

as well as between the cross-sectional area of the branch, *A*_total_, and its lumen, *A*_lumen_,

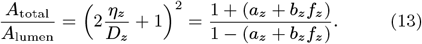

Similarly, at the outer surface of the epithelium,

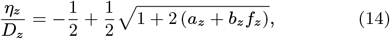

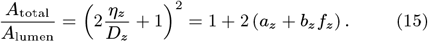

These relationships depend on the same two dimensionless biophysical parameters, the pressure difference between the tree opening and the exterior of the bronchial tube, *p*_base_ − *p*^out^, and the shear stress, *τ*, both relative to the sum of hoop stress and pressure on the exterior of the bronchial tube, *σ*_*θ,z*_ + *p*^out^:

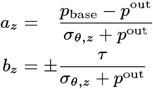

*σ*_*θ,z*_ is expected to change within the epithelial wall, while the other parameters are the same everywhere within it. Similar relationships can also be obtained with the thin-walled cylinder approximation, but the hoop stress change within the epithelial wall is then ignored (Supplementary Text). For accuracy, we will use the thick-walled cylinder approximation going forward.

Both Eq. 12 and Eq. 14 can fit the data with uniform hoop and axial stress in all branches, i.e., *a*_*z*_ = *a* and *b*_*z*_ = *b*, but the equations would predict slightly different thickness-diameter relationships, and the fit on the luminal side is statistically favoured (Fig. 5f, Table S1). The inferred biophysical parameter *b* increases slightly over developmental time, while *a* decreases (Fig. 5f, inset, Table S1), which is consistent with a reported decrease in amniotic fluid pressure during gestation [54], though our estimate is based on a single sample per developmental time point. Assuming uniform material properties of the epithelial walls, uniform hoop and axial stress implies that all cells experience the same strain, i.e., the same amount of relative deformation. The observed diameter-thickness relationship can thus emerge if epithelial cells remodel their shape between a flat squamous and a highly elongated pseudostratified configuration to maintain a preferred level of strain.

We conclude that the bronchial tree, despite its asymmetry, is organised such that the shear stress, the pressure drop, as well as the axial and hoop stress equalise in all branches. An increase in the volume flow via an increase in the tip number or in the rate of fluid secretion at the tips would result in circumferential growth, while a loss in flow due to destruction of alveoli would lead to circumferential shrinkage. Once branching morphogenesis is complete at the end of the pseudo-glandular stage, further lung growth can be expected to result primarily in branch lengthening. The increase in thoratic space will permit additional alveologenesis that results in additional flow, which in turn results in tube widening. As both the branch length and the inner diameter cubed change in parallel to the number of tips, the pressure drop per branch will remain unchanged. The change in diameter will nonetheless result in a change in wall thickness to maintain the same hoop stress. Considering that the relationships that we uncover in the embryonic mouse lung (Fig. 5) also apply to the adult mouse and human lung (Fig. 1), the lung likely grows and remodels by the same physical principles throughout adolescent and adult life.

### Reorganisation of the juvenile canine bronchial tree after pneumonectomy

To follow adult lung remodelling over time, we re-analysed data from two litter-matched foxhounds that underwent removal of the right lung (PNX) or right thoracotomy without lung resection (SHAM) at 2 months of age, and that were subsequently followed until somatic maturity at 12 months of age [53]. Vigorous compensatory growth of alveolar septal tissue and respiratory bronchioles completely normalises lung volume, gas exchange, and maximal oxygen uptake [53]. Spiral CT scans, performed at 6 months of age (4 months after surgery) and in three animals of each group at 12 months of age (10 months after surgery), show that the lung tree post-PNX is structurally and functionally distinct from the control (Fig. 6a). The chest cavity changes substantially after treatment and the quality and quantity of remodelling depends on the type of recession [55]. All the remaining lobes increase in volume, while heart and mediastinal structures are shifted to the right side of the chest. Portions of the left cranial and caudal lobe moved across the midline, wrapping above, below, and around the heart to fill the empty space in the right hemithorax. Given that the pulmonary viscous resistance and ventilatory power requirements measured at any given ventilation at rest and during exercise are approximately 2–2.5 times those in normal dogs, while elasticity is increased in PNX lungs [53], we wondered whether the lung post-PNX still follows the established biophysical relationships. We can segment individual branches up to generation 11, but segmentation of all sister branches is possible only up to generation 5–6. Given the increasing inaccuracy, we limited our analysis to the first seven generations.

**Figure 6:**
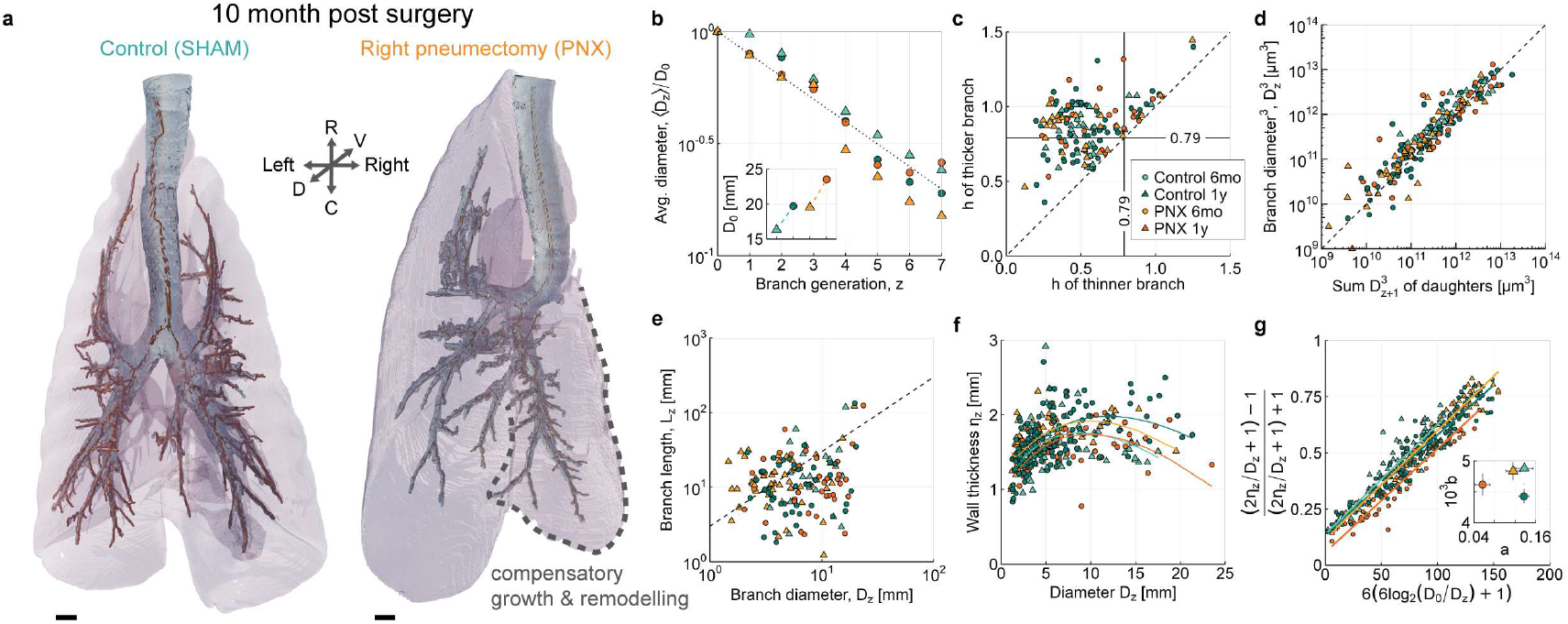
Reorganisation of the adult canine bronchial tree after pneumonectomy. **a**, 3D rendering of airways of control foxhound bronchial trees and 10 months after pneumonectomy at 12 months of age. Scale bars: 10 mm. R: rostral, C: caudal, D: dorsal, V: ventral. **b**, Average inner branch diameters against branch generation *z*, starting at the trachea (*z* = 0) in adult canine control (SHAM) (teal) and treated lungs (orange). Dashed line represents the Hess–Murray law. Insets: Trachea diameter in each sample. **c**, Homothety ratio *h* of the inner branch diameters of sister branches in adult canine control (teal) and treated lungs (orange). **d**,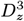 of parent branch vs. 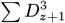 of their daughter branches in control (teal) and treated lungs (orange). **e**, Branch length-to-diameter ratio. Dotted line indicates *L*_*z*_ = 3*D*_*z*_. **f**, Airway wall thickness versus diameter in SHAM (control) and treated (PNX) lungs. **g**, Data from **f** transformed to infer biophysical parameters (inset, error bars are s.e.) by fitting Eq. 12 (lines in **f**,**g**). All analyses are based on *n* = 1 image per time point and condition. CT data re-analysed from [53].

Consistent with previous reports [53], we confirm a ≈ 20% dilation of the trachea (Fig. 6b, inset, based on a single sample, *n* = 1). Canine lungs have been suggested to be particularly asymmetric [56], but we observe similar differences between sister branches (Fig. 6c Fig. S6b) as in adult mice (Fig. 1e). The average inner branch diameters follow the Hess–Murray law, in particular after pneumonectomy (Fig. 6b, Fig. S6a,c,d), while the individual inner branch diameters follow Murray’s law (Fig. 6d). Even though the individual branch length-to-diameter ratios deviate substantially from 3 (Fig. 6e), suggesting different pressure drops across branches (Eq. 4), the relationship between the wall thickness and the inner branch diameter (Fig. 6f) can again be fitted well by Eq. 12, using the same parameter values for *a, b* for all branches (Fig. 6g). The impact of a vertical pressure gradient oriented from the apex of the lung to its base can be incorporate in the model, but has a minimal impact on the fitted parameters (Table S2). While *a* remains unchanged, *b* decreases slightly with age in the control lung. Pneumonectomy results in smaller *a*, but a smaller decrease in *b* with age (Fig. 6g, inset).

We conclude that lungs grow and remodel by the same physical principles throughout adolescent and adult life. This explains how the lung can maintain its energy-efficient architecture as the organism grows. One would, however, also expect remodelling in disease.

### Healthy and COPD-affected lungs differ in biophysical parameters

Chronic obstructive pulmonary disease (COPD), the third leading cause of death worldwide [58], is a chronic inflammatory lung disease that causes obstructed airflow from the lungs [59]. The two most common conditions that contribute to COPD are emphysema, in which the alveoli are destroyed as a result of damaging exposure to cigarette smoke and other irritating gases and particulate matter, and chronic bronchitis, an inflammation of the lining of the bronchial tubes. Early stages of COPD are often difficult to diagnose, not least because the resolution of current imaging technologies is insufficient to detect the destruction in individual alveoli or bronchi.

We wondered whether we would detect changes in the organisation of the upper pulmonary tree in COPD patients. We analysed 42 chest CT scans from healthy humans and 796 CT scans from 533 different COPD patients [57]. Given the resolution limit of 1 mm per voxel, it was not possible to extract the branch diameters, lengths, and wall thicknesses beyond generation 12 in controls and beyond generation 15 in COPD patients (Fig. 7a) of the up to 26 branch generations in human lungs [4, 5]. We find the cubed diameters of parent branches to strongly correlate with the sums of the daughter branches, which is indicative of equal shear stress in all branches (Fig. 7b), as argued before for the adult human lung [43]. Unlike in the mouse embryo (Fig. 5e), the wall thickness in human bronchial trees appears constant for generation 0–3 (Fig. 7c,d) [60], but due to limitations of the CT method, the thickness quantification in generations *z* = 0–2 is unreliable [57]. In the first four airway generations, flow is likely turbulent in human lungs [61–63]. Poiseuille flow in a pipe is associated with the minimum pressure drop for a given flow rate; thus all other forms of flow necessarily require a greater pressure drop for the same flow rate [62]. However, from generation 4 onward, the expected error in the estimated pressure drop becomes fairly small [62] and the average length-to-diameter ratio becomes linearly related [64]. We will therefore continue to use this simple approximation, but limit our analysis to *z* g 4. Using Eq. 12, we find that the airway thickness increases with the airway diameter in a way that the hoop and axial stresses remain constant in all branches (Fig. 7e). Remarkably, the relative biophysical parameters *a* and *b* lie within a similar range as in the mouse embryo and the canine lungs, despite the very different size of the adult human lungs (Fig. 7e,f, Fig. S7). The impact of a vertical pressure gradient oriented from the apex of the lung to its base can be incorporate in the model, but has a minimal impact on the fitted parameters (Table S3, Fig. S8). We could not detect any statistically significant correlation of the parameters with age (Fig. 7f). Males exhibit significantly smaller *a* values (*p <* 0.05) and larger *b*/*a* ratios compared to females, while *b* is increased only in male COPD patients relative to female patients (Fig. S9). In COPD lungs, *b* and *b*/*a* are significantly larger in both sexes. Higher *b* and *b*/*a* values are consistent with higher shear stress in forced expiration [43, 65]. Consistent with spatial differences in COPD lungs, *a* and *b* deviate significantly more between the left and right lung in COPD patients than in healthy controls, in particular in those lungs with the highest *a, b* values (Fig. 7g,h).

**Figure 7:**
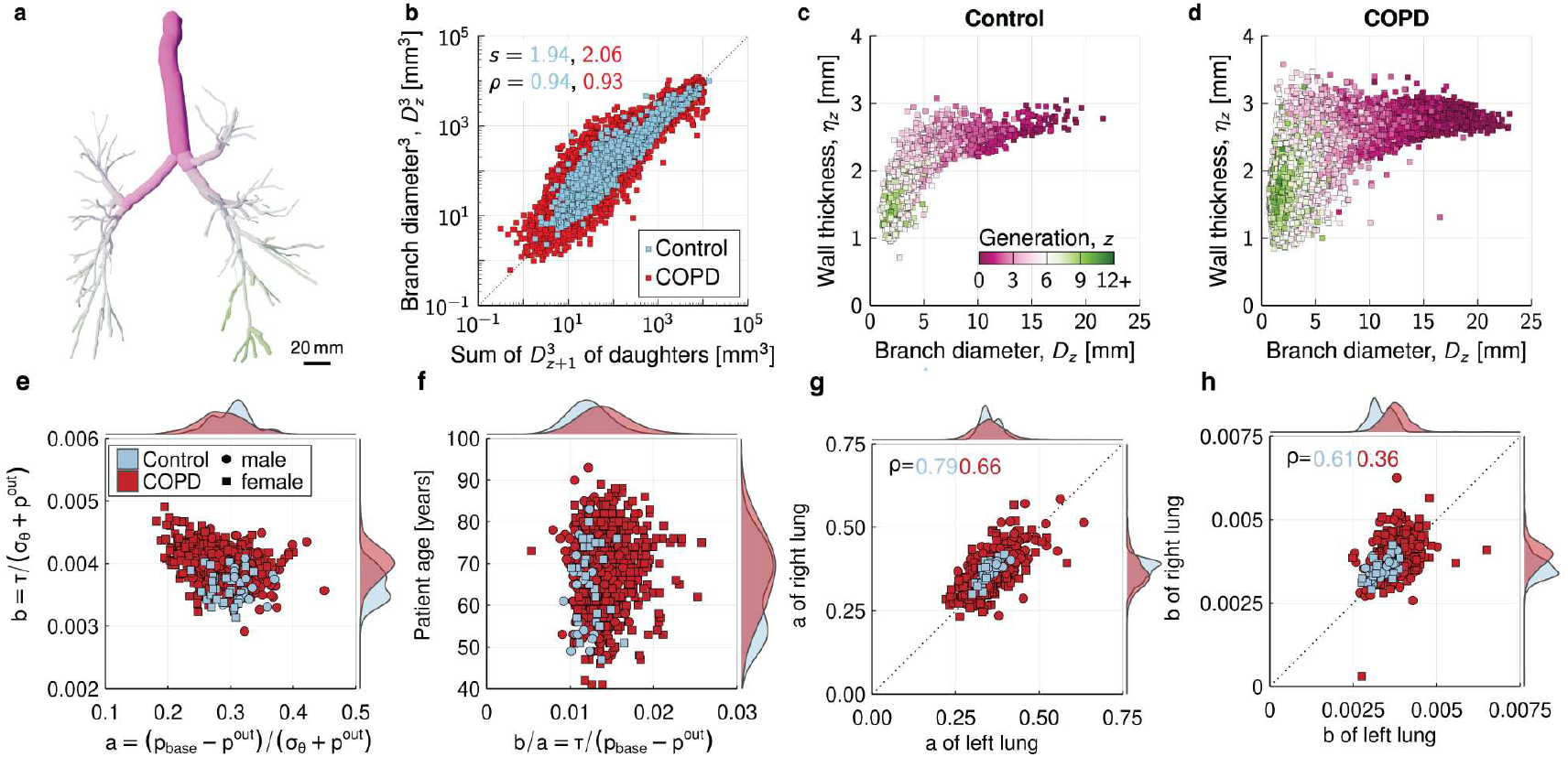
Difference in the relationship between airway diameter and wall thickness between healthy and COPD patients. **a**, 3D rendering of the segmented airways of an 80-year-old woman with COPD, including branches down to generation 12. **b**,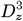 of parent branch vs. 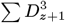 of their daughter branches in adult human control (light blue) and COPD lungs (red). Variability is quantified by the standard deviation 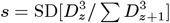. **c**,**d**, Airway wall thickness versus diameter in control (c)and COPD **(d)** lungs. **e–h**, Inferred biophysical parameters in entire lungs **(e**,**f)**, or in the left or right lung **(g**,**h)** of control and COPD patients. Parameters were obtained by fitting Eq. 12 to the data of individual lungs in **(c**,**d)**. Fitted distributions are shown on the plot boundaries. *ρ* are Pearson correlation coefficients. Total number of lung images of control: *n* = 42 **(b–f)**, *n* = 37 **(g**,**h)**, an d COPD patients: *n* = 796 **(b–f)**, *n* = 720 **(g**,**h)**. CT data re-analysed from [57].

## Discussion

### The Developing Lung: Fractalization and Asymmetry

For over a century, the lung’s fractal structure has intrigued scientists, yet the processes underlying its development, in time for the first breath at birth, have remained elusive. Seminal work by Hess, Murray, and Weibel demonstrated that the bronchial tree adheres to geometric principles for energy-efficient ventilation [4, 6, 7]. However, how the lung self-organizes into this architecture and whether fractalisation is gradual or sudden were unresolved questions. Adding to this complexity, mammalian lungs exhibit local deviations from global optimization rules. Are these deviations merely reflections of natural variability and imprecise development, or do they emerge from physical principles with specific functional purposes?

### Filling the Observational Gap in Lung Morphogenesis

The intermediate stages of lung development, when the fractal structure emerges, have been largely inaccessible. Previous studies focused on early developmental stages using 3D imaging, 2D cultures and perturbation analyses [27, 30, 66, 67] or adult lungs using micro-CT and casting techniques [68–70]. We bridge this gap with a 4D dataset spanning the pseudo-glandular phase of lung development and introduce the computational tool SkelePlex to analyse the network morphologies.

### A new tool for 3D image-based biological network analysis

SkelePlex is a versatile, open-source tool for extracting morphological features from complex networks, offering researchers a powerful resource for studying pulmonary systems and beyond. By integrating advanced 3D imaging with neural network analysis, SkelePlex enabled us to uncover the organ-wide dynamics of tube diameter and wall thickness remodelling that drive the formation of the lung’s fractal, energy-efficient architecture. Designed for broad applicability, SkelePlex supports diverse imaging modalities and network structures, making it a versatile resource for studying biological and other systems.

### Biophysical Forces Driving Fractalization

Our image-driven biophysical analysis revealed that lung fractalisation during organogenesis is a dynamic, gradual process driven by fluid-structure interaction. The branch wall thicknesses are established concomitantly. We found that wall thickness and diameter are correlated in a way that maintains uniform hoop stress in the epithelial wall, identifying a new biomechanical principle underlying lung morphogenesis. This uniform hoop stress is intrinsically linked to a uniform luminal pressure drop [10] and a uniform shear stress exerted on the epithelial wall by intrapulmonary flow [42, 43]. Together, these forces establish a mechanically optimized architecture that supports efficient air flow and gas exchange.

Notably, these principles extend beyond development to govern the lung’s response to perturbations after birth. In conditions such as partial lung regrowth following pneumonectomy or widespread tissue damage in COPD, the bronchial tree dynamically remodels to restore uniform mechanical conditions. This persistence of developmental mechanisms into adulthood enables the lung to adapt to changes in mechanical stresses. Such adaptive plasticity ensures that the conducting airways reorganize to maintain an optimal, energy-efficient structure. These findings demonstrate that lung morphology adapts dynamically to preserve uniformity in key mechanical forces—shear stress, hoop stress, and pressure drops—across the bronchial tree, ensuring the organ’s functional resilience under both developmental and pathological conditions.

### Biophysical Metrics as Biomarkers

We identified dimensionless, interpretable, spatially-uniform bio-physical parameters

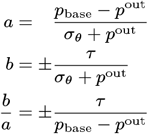

that link tissue stress levels to fluid pressure, enabling a quantitative description of lung architecture across species and developmental stages. These parameters not only highlight the simple, uniform principles governing lung organization but also hold potential as biomarkers for tracking and diagnosing changes in lung architecture over time. For example, airway obstructions are predicted to alter stress and shear forces, triggering compensatory remodelling, reflected in the parameters. Longitudinal studies with higher temporal resolution will be critical to validate these predictions.

### Linking mesoscopic biophysical principles to cellular mechanisms

Understanding how mechanotransduction converts biophysical cues—such as shear and hoop stress—into cellular responses is essential for linking mesoscopic principles to cellular mechanisms, and for elucidating their roles in lung morphogenesis and pathology. Recent studies have highlighted the involvement of mechanosensitive ion channels, such as TRP, Piezo, and TREK channels, in lung biology [71]. For instance, TRP channels, including TRPV4, play a critical role in airway remodeling in diseases like asthma and COPD [72, 73], while Piezo1 mediates epithelial responses to mechanical stretch [74, 75]. These channels are critical in sensing and responding to mechanical stress, influencing growth and remodeling. Our biophysical model provides a framework to guide this research by predicting the expected branch morphology based on geometric principles, and offering clear hypotheses about how cellular responses should vary in response to deviations from these expectations.

### Conservation across biological networks

Since Murray’s seminal work, various biological networks, such as blood vessels, respiratory systems, nutrient transport in Amoebozoa, and xylem in plants, have been shown to follow the Hess-Murray law [4, 76–79], though deviations are observed in others [80, 81], likely reflecting deviations from Newtonian fluid behaviour and other key assumptions [82]. It is thus plausible that similar mechanosensation mechanisms are common in transporting networks. In microvascular networks, wall and shear stress regulate structural adaptation over time [80, 83]. Similarly, *Physarum polycephalum* responds to shear stress, adjusting its cytoplasmic network [34, 35]. Unlike the lung, these systems rely on expansion followed by pruning and branch number reduction [34, 84]. While the core mechanisms are broadly shared, their implementation varies, reflecting adaptations to specific functional demands. As such, comparisons across systems must be done carefully [85–88].

### Concluding remarks

Our study demonstrates that the lung’s fractal structure emerges through simple, uniform biophysical principles that govern both development and adaptation. By bridging scales from biophysical principles to organ-level morphology, we provide a foundation for future research exploring how mechanical forces shape development and adaptation across systems. By linking biophysical forces to network morphology, we provide a framework for studying mechanically sensitive tissues and uncovering biomarkers for disease. These insights not only advance our understanding of lung biology but also highlight the broader applicability of these principles across biological systems, offering new avenues for research and therapeutic strategies.

## Limitations of the study

### Experimental accessibility

Although natural and surgical perturbations in mice and other model organisms support the assumptions and predictions of our proposed mechanisms, the experimental challenges associated with later embryonic stages limit comprehensive validation of causal relationships [37, 41, 51, 89]. Traditional lung explant cultures, which are well-established for the early stages of lung development until E12.5 [90], are inadequate to investigate the impact of fluid flow and pressure perturbations on branch remodelling in later lung developmental stages that depend on active blood supply, pressure gradients, mechanical forces and chest integrity. While grafting lungs under the renal capsule extends culture duration, it prevents real-time measurements during the culture process [91]. Additionally, by E15.5, mouse embryonic lungs can reach several centimeters in size and become too thick and opaque for effective live tissue imaging using conventional light microscopy. While advanced two-photon microscopy offers improved deep tissue imaging, it is currently limited to depths of 1 mm and field-of-view sizes up to 2 mm^2^, which are insufficient to capture the architecture of entire murine embryonic lungs [92]. Therefore, significant advancements in both organ culture methods and imaging technologies are required to enable effective probing and manipulation of fluid flow during the late pseudo-glandular stage in mice.

### Theoretical model

Our theoretical model assumes laminar flow, uniform material properties, and linear elasticity to derive biophysical principles. While these simplifications adequately describe the available data, they ignore several important factors studied previously [82, 93–96]. The dynamic viscosity of air varies with temperature: As air warms up from ambient (25°C) to body temperature (37°C) during breathing, its dynamic viscosity—and consequently the shear stress (Eq. 3)—increases by approximately 3%. During fetal development in homeothermic animals, the temperature of amniotic fluid is expected to remain constant. As such, we expect the effects of temperature variation to be negligible, but we cannot exclude that other effects result in spatial difference in the dynamic viscosity. We further assume the stress-strain relationship to be independent of the axial position in the lung. In upright position, there is a gravity-based pre-load of the lung tissue, compressing the lung at the base while stretching it at the apex [97]. During development, however, a frequently rotating fetus should not be exposed to this anisotropy.

Deviations from cylindrical tube shapes—such as bends, rough surfaces, and non-circular cross-sections—impact the flow profile [98].

### Pathological progression over time

The existing human patient dataset [57] does not allow for longitudinal tracking of disease progression in COPD patients. Future studies monitoring the parameters *a* and *b* over time could provide valuable insights into the progression of pulmonary pathologies. Such longitudinal data might inspire novel drug design strategies and offer metrics for evaluating treatment success.

## Materials and Methods

### Mouse strains

We used embryonic lungs from mice homozygous for the Rosa26mTmG and heterozygous for the ShhGFP-Cre allele (ShhGFP^Cre/+^; Rosa26mTmG) [99]. The double-fluorescent SHH-controlled Cre reporter mouse expresses membrane-targeted tandem dimer Tomato (mT) before Cre-mediated excision and membrane-targeted green fluorescent protein (mG) after excision [100]. As a result, the epithelial cell membranes in the lungs are labelled by GFP, while all adjacent mesenchymal tissue is labelled by tdTomato.

### Animal housing and husbandry conditions

All mice included in the study exhibit a healthy phenotype and have not been utilized in any prior experiments. Housing and husbandry were performed following the guidelines and with the protocols of the veterinary office of the Canton Basel-Stadt, Switzerland.

### Lung collection

The developmental stage of mouse embryos was identified by performing daily vaginal plug checks during breeding. Embryonic lungs of the desired developmental stage were carefully dissected using fine forceps and tungsten needles in cold phosphate-buffered saline (PBS), and then collected in a Petri dish. The lungs were subsequently fixed in 4% paraformaldehyde (PFA) for a duration of 2–6 hours, depending on their size. Fixation was concluded by subjecting the specimens to five 10-minute washes, followed by an additional overnight wash.

### Optical clearing and immunoflurescence

Whole-mount tissue clearing of dissected lungs was performed with the Clear Unobstructed Brain/Body Imaging Cocktails and Computational Analysis (CUBIC) protocol, as previously specified [99, 101]. Clearing times in reagents for decolouring, delipidation, permeation (CUBIC-1) and refractive index (RI) matching (CUBIC-2) were adjusted to maximize clearing efficiency and minimize quenching. The samples were incubated in 50 % CUBIC-1 (CUBIC-1:H_2_O=1:1) for 2 days, and in 100 % CUBIC-1 for 3–14 days until they became transparent. All samples were subsequently washed several times in PBS. The immunostaining was performed after the CUBIC-1 clearing steps. All samples were then treated with 50% CUBIC-2 (CUBIC-2:PBS=1:1) for 2–5 days. Lastly, incubation in 100% CUBIC-2 was continued until the desired transparency was achieved, 3–5 weeks, with CUBIC-2 being refreshed every 2 days. All CUBIC steps were performed on a shaker at room temperature. Cleared samples were embedded in 2% low-melting-point (LMP) solid agarose cylinders, and immersed in CUBIC-2 overnight.

To enhance the epithelial fluorescent signal, we stained for Keratin 8 (Krt8), using anti-Krt8 (TROMA-I) rat monoclonal antibody (Hybridoma Bank). Endogenous GFP signal was amplified by performing immunostaining with Anti-Green Fluorescent Protein (GFP) chicken polyclonal primary antibody from Aves Labs (1:200). After CUBIC-1 treatment lungs were washed four times for 15 min in PBS. Lung samples were blocked overnight in 4% bovine serum albumin supplemented with 0.3% Triton-X. The next day primary antibody was added in fresh blocking solution in 1:200 dilution for 48 hours at 4°C. Samples were then washed with PBS 3 times for 15 min and once overnight. Then samples were incubated with a fluorescently conjugated secondary antibody diluted 1:500 overnight (Alexa Fluor 555: Invitrogen; A-31572) and washed with PBS, following CUBIC-2 optical clearing.

### Light-sheet imaging

3D images of E10.5–E13.5 lungs were previously published [30]. We imaged the E14.5 lungs with a Zeiss Lightsheet Z.1 microscope using a Zeiss EC Plan-Neofluar 5x/0,16 M27(420330-9901-000) objective. Owing to the size of the E15.5 and E16.5 lungs, we imaged them using a mesoSPIM v5 light-sheet microscope with a macro-zoom system (Olympus MVX-10) objective [102]. Given the size of the samples, image volumes were acquired in tiles and stitched using BigStitcher v1.2.11 [103]. To enhance local contrast, the images were deconvoluted using Huygens Professional v19.04 (Scientific Volume Imaging, Netherlands). Images were downsampled to generate isotropic voxel sizes and the local contrast was enhanced using contrast limited adaptive histogram normalisation.

### Segmentation of mouse embryonic bronchial tree

We segmented the lung branches using a 3D U-Net. We used napari [104] and APOC [105] to densely label 512^3^ voxel patches of the images. Using these annotated patches, we then trained a 3D U-Net to segment the lung branches from the light-sheet images. Clearing large embryonic lung tissue is challenging. We therefore used two different samples for the E16.5 time point. To reconstruct the complete E16.5 lung graph, we performed an extended clearing and utilized the auto-fluorescent contrast between tissue and bronchial lumen to segment the luminal volume as the endogenous GFP signal was poorly preserved. To quantify the epithelial wall thickness, we shortened the clearing process to preserve the endogenous GFP signal; this did not provide a clear image of the entire lung and only allowed us to reconstruct the graph partially up to generation 11.

### Segmentation of canine chest CT images

The bronchial tree of the canine chest CT images was segmented using the Dragonfly Software (v2024.1) [106]. The resulting label images were manually refined and used to train another 3D U-Net to be more generally applicable for similar datasets.

### Development of SkelePlex

To skeletonise the lung tree from light-sheet microscopy images, we trained a multiscale 3D U-Net to predict the skeleton from the segmented lung branches. We generated training data by creating synthetic segmentations of lung trees with varying branch lengths, branch diameters, and branching angles. As a pre-processing step, we performed a Euclidean distance transform on the segmentations, in which each voxel holds the distance to the nearest background voxel, normalised by the maximum distance in the surrounding 5^3^ voxel neighbourhood. We implemented a 3D multiscale U-Net (a variant of the U-Net architecture with an additional output at each decoder layer) in PyTorch and trained it to predict the skeleton from the normalised distance image. As an auxiliary task during training, the network predicted the skeleton at each scale of the decoder.

### Training of SkelePlex skeletonisation network

We trained the SkelePlex skeletonisation network using simulated lung branch segmentations. We generated realistic lung branch segmentations by first creating a fractal skeleton graph and then expanding it into a densely-labelled segmentation image. We additionally randomly displaced the surface of the resulting simulated lung branch segmentations to emulate the bumpy surface observed in the real images. In generating the training data, we randomly varied the branch angle (10–130°) and branch diameters (8–52 voxels). We divided the data with a 90:10 training-validation split and trained the skeletonisation model using the Adam optimiser with a learning rate of 0.0004. We computed the loss as the sum of the smooth *L*_1_ losses [107] for each scale output of the multiscale 3D U-Net. We augmented the training data with random flips, random 90° rotations, and random affine transformations (rotation: ±1.5 radians, translation: ±20 voxels, scale 1–1.1).

### Analysis of bronchial trees

To quantify the architecture of the developing lung trees, we developed a pipeline (SkelePlex) to construct graph representations of the bronchial tree as well as to measure branch lengths and diameters from 3D light-sheet microscopy images. First, we segmented the lung branches using a 3D U-Net. Then, we skeletonised the segmentation using the skeletonisation network.

Using the resulting skeleton, we constructed a graph where the branch points of the tree are nodes and the branches are edges. We visualized and curated the lung branch trees using napari [104]. To measure the lengths, diameters and wall thicknesses of the branches, we fit splines to the branches. With the branch splines, we directly measured branch lengths, diameters and wall thickness in orthogonal planes using the morphometrics package [108]. Diameter and wall thicknesses are computed from segmented areas in these othogonal image slices.

## Code Availability

The source code is released under the 3-clause BSD license. It is available as a public git repository at https://git.bsse.ethz.ch/iber/Publications/2025_mederacke_fractal_lung.

## Acknowledgements

We thank Noah Grodzinski and Patrick Jenny for valuable discussions. Part of imaging was performed with equipment maintained by the Center for Microscopy and Image Analysis, University of Zurich.

## Author Contributions

MM, ND, LS obtained the embryonic data; KY, MM developed SkelePlex; CH, MD provided the canine CT data; JS, TW, MP, JB provided the quantified human CT data; DI developed the theory and supervised the project; MM, RV analysed the data. DI, MM, KY, RV, ND wrote the manuscript; RV, MM, KY prepared the figures. All authors reviewed and approved the manuscript.

## SUPPLEMENTARY MATERIALS

Supplementary Text

Figures S1 to S4

Tables S1, S3, S2

References

## Supplementary Text

### Impact of variable tube diameter on shear stress

Variations in the tube diameter can impact our estimate of the shear stress. The average shear stress in case of a variable tube diameter is given by

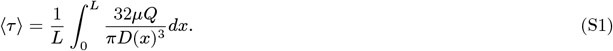

Here, *L* is the length of the branch and *D*(*x*) is the diameter along the branch. The fold-increase, *F*, relative to a uniform tube diameter is

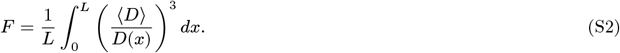

The observed increase in shear stress due to variable diameters of a single branch are small (Fig. S4).

### Thin-walled cylinder

If the wall thickness is considerably smaller than the tube diameter (*η* ≪ *D*), the thin-walled cylinder approximation can be used. For a thin-walled cylinder, the hoop stress *σ*_*θ*_ is approximately two times the axial stress *σ*_*L*_ and is given by:

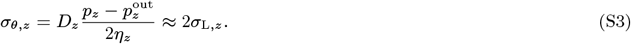

In combination, Eqs. S3 and 10 yield a simple relationship between wall thickness and branch diameter:

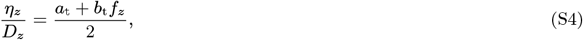

where

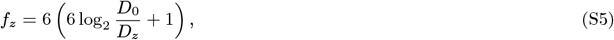

with two dimensionless biophysical parameters:

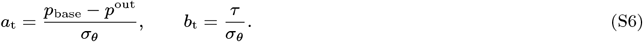

### Thick-walled cylinder

When the wall thickness and lumenal diameter are of similar dimensions, as is the case in the bronchial tree, the thick-walled cylinder approximation is more appropriate.

#### Axial Stress

In case of a closed-ended cylinder with thick walls, the axial stress in longitudinal direction in the tube wall is given as

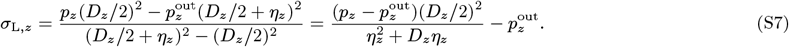

This can be written as

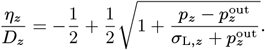

The second solution of the quadratic equation is unphysiological as *η*_*z*_/*D*_*z*_ *>* 0. Using Eq. 10, this can be rewritten as

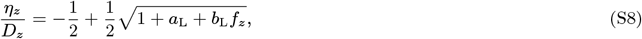

which has two dimensionless biophysical parameters,

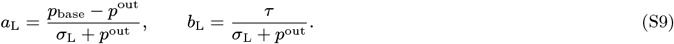

Note that *f*_*z*_ is the same as in the thin-walled approximation (Eq. S5).

#### Hoop Stress

The hoop stress at distance *d* ∈ [0, *η*_*z*_] from the inner wall surface towards the outer surface of an open or closed-ended cylinder is given by

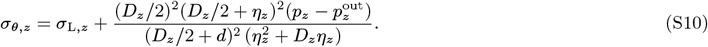

On the lumenal side, where *d* = 0,

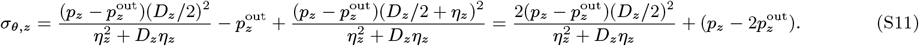

This can be written as

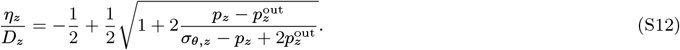

The second solution of the quadratic equation is unphysiological as it violates *η*_*z*_/*D*_*z*_ ≥ 0. Using Eqs. 10 and 11, this can be rewritten as

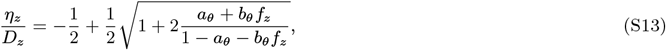

which has two dimensionless biophysical parameters,

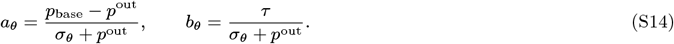

To facilitate parameter inference from the data, this can be rearranged to a linear relationship in *a*_*θ*_ and *b*_*θ*_ :

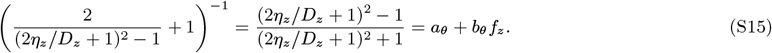

At the outer surface, where *d* = *η*_*z*_, we have

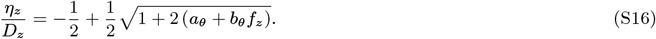

This can be rearranged to

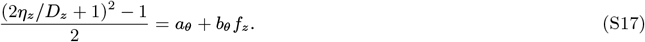

Notice that *b*_*θ*_/*a*_*θ*_ = *b*_L_/*a*_L_ = *τ/*(*p*_base_ − *p*^out^) = *b*_t_/*a*_t_, as in the thin-walled approximation. For each branch, only *σ*_*θ*_ can differ between the lumenal side and the outside. Accordingly, the fractions *b*/*a* are expected to be equal on the lumenal and outer surface 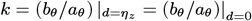, such that Eq. S16 becomes

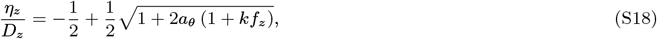

or after rearrangement,

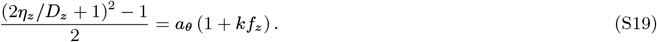

Fitted values for the human lung are provided in Table S3.

#### Radial Stress

The radial stress at distance *d* ∈ [0, *η*_*z*_] from the inner wall surface towards the outer surface is given by

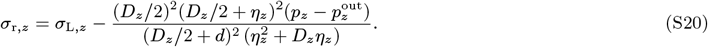

On the lumenal side, where *d* = 0,

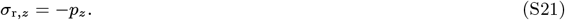

At the outer surface, where *d* = *η*_*z*_,

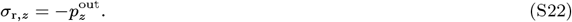

### Impact of a vertical pleural pressure gradient

In the adult lung, a pressure gradient has been reported in the intrapleural space, i.e., in the pressure that is measured in the space between visceral and parietal pleurae [48]. Using different techniques, in head-up dogs, the gradient has been estimated as 20–90 Pa/cm vertical distance [48]. Assuming that this gradient is indeed linear, we replace *p*^out^ by *p*^out^(1 + *gG*), where *G* is the vertical distance between the lower end of the trachea and the branch element.

#### Thin-walled cylinder

Taking into account the pleural pressure gradient, the equation for the thin-walled cylinder (Eq. S4) becomes

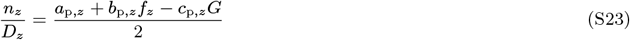

with three parameters

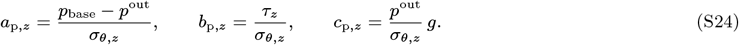

#### Thick-walled cylinder

Taking into account the pleural pressure gradient, the equations for the thick-walled cylinder (Eqs. 12 and Eq. 14) becomes on the lumenal surface

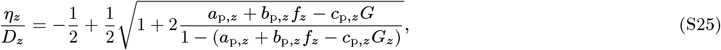

and on the outer surface

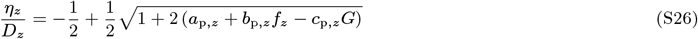

with the same three parameters as in Eq. S24. When fitting the human lung data with Eq. S25, we obtain *b*/*a* ratios similar to the ratios obtained with the alternative models (Table S3, Fig. S8). The inclusion of a vertical pleural pressure gradient thus does not substantially alter our inferred biophysical parameters.

### Relation of fitted biophysical parameters to quantitative measurements

Our models relate the hoop and shear stresses to the hydrostatic pressure in the branch lumen and on the outer epithelial surface. In the following, we relate our inferred relative parameters to measured values.

The hoop stress experienced by lung epithelial cells has been proposed to be about 3 cm H_2_O ≈ 300 Pa in normal breathing and about 30 cm H_2_O ≈ 3000 Pa under smooth muscle mediated bronchoconstriction [109]. Others inferred much higher hoop stresses of about 5000 Pa [110] based on measurements of intact airways segments of generation 10 and below [111]. The stress must then, however, mainly be absorbed by the ECM, considering that lung epithelial cells have a Young’s modulus in the order of 500 Pa [112]. Measurements suggest a strain of 15–20% for physiological pressure levels [112].

Before inspiration occurs, the pressure in the pleural space is negative (≈ −500 Pa), which keeps the lungs inflated. As the diaphragm contracts, the volume of the thoracic space increases and pressure in the pleural space decreases (it becomes more negative in relation to atmospheric pressure), i.e., the pressure in the pleural space drops from about −500 Pa to about −1000 Pa at the end of inspiration. Accordingly, transmural pressure is in the range of 500–1000 Pa. However, *p*^out^, the pressure on the outer epithelial surface, may be different from the pleural pressure. Indeed, if we set *p*^out^ to the expected pleural pressure, then *a* = (*p*_base_ − *p*^out^)/*σ*_*θ,z*_ ∈ [500/300, 1000/300] ≈ [1.5, 3], which is 5–10 times higher than our estimate. We can obtain consistent values for *a* if we assume that *p*^out^ ≈ −100 Pa, suggesting that there is a pressure gradient in the parenchyme.

Airway epithelial cells are exposed to luminal wall shear stress (frictional force per surface area) generated by airflow, which at rest is 0.5–3 dynes/cm^2^ = 0.05–0.3 Pa [113–115]. Shear forces may be a lot higher during exercise and may increase to nearly 1,700 dynes/cm^2^ = 170 Pa with cough and bronchospasm [113]. Together, the approximation for the thin cylinder, *b*_t_ = *τ/σ*_*θ*_ ∈ [0.16, 1] × 10 ^−3^, is 10-times smaller for normal breathing than inferred, or even lower using the inferred values. Under exertion, *b*_t_ ought to further increase as *τ* appears to increase more than *σ*_*θ*_. If we now consider *b*_*θ*_ = *τ/*(*σ*_*θ*_ + *p*^out^) from the thick-walled approximation, we have *b*_*θ*_ ≈ 0.2 Pa/(300 Pa − 1000 Pa) ≈ −3 × 10^−4^, which is about 10 times smaller than our inferred values. With *p*^out^ ≈ −100 Pa, the inferred *b*_*θ*_ would still be 3-times smaller; *p*^out^ ≈ −240 Pa would be needed for a perfect match.

Using different techniques and different species, the gradient in the pleural pressure has been estimated as −20 to −90 Pa per centimetre vertical distance [48, 49]. In our model, the vertical pressure gradient is given by *p*^out^*g* = *cσ*_*θ*_ = (*c/a*)(*p*_base_ − *p*^out^). For our fitted values *c/a* = (−1 ± 60) × 10^−4^/cm for control and *c/a* = (−2 ± 70) × 10 ^−4^/cm for COPD patients and with *p*_base_ − *p*^out^ ≈ 100 Pa, we obtain a pleural pressure gradient along the outer epithelial surface that is almost a 1000-times shallower than the reported pleural pressure gradient, with a large variation between samples. This may be due to parenchymal material effects that dampen the pleural pressure gradient.

**Fig. S1:**
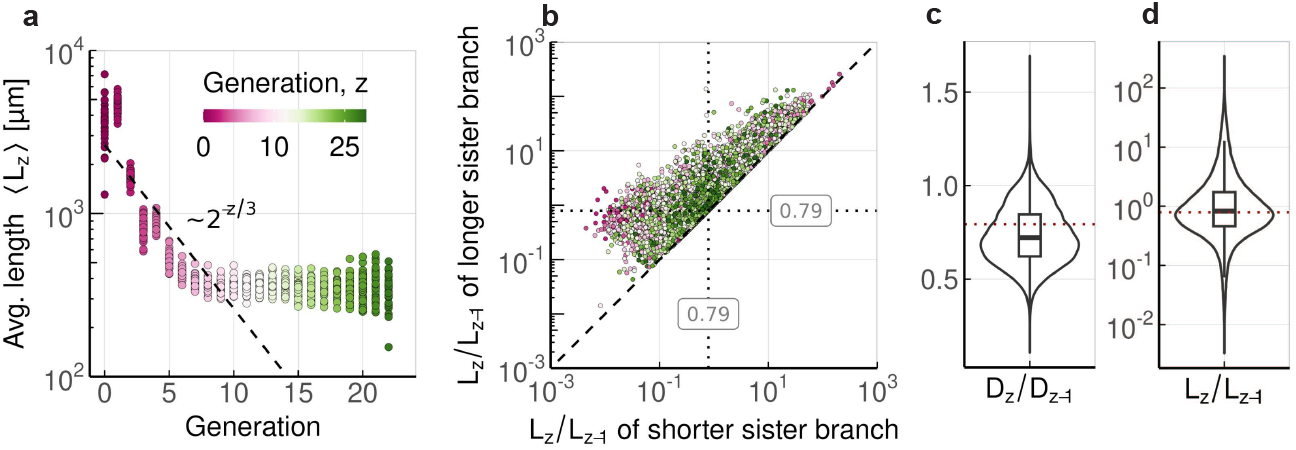
Morphometrics of the lungs in adult mice. **a**, Average branch lengths against the branch generation. Dashed line represents the Hess–Murray law. **b**, Homothety ratio of the branch lengths of sister branches. **c**,**d** Variability of the homothety ratio of the inner branch diameters **(c)** and branch length **(d)**; *h* = 2^−1/3^(dotted lines) for comparison. Data from [1], *n* = 34 mouse lungs.

**Fig. S2:**
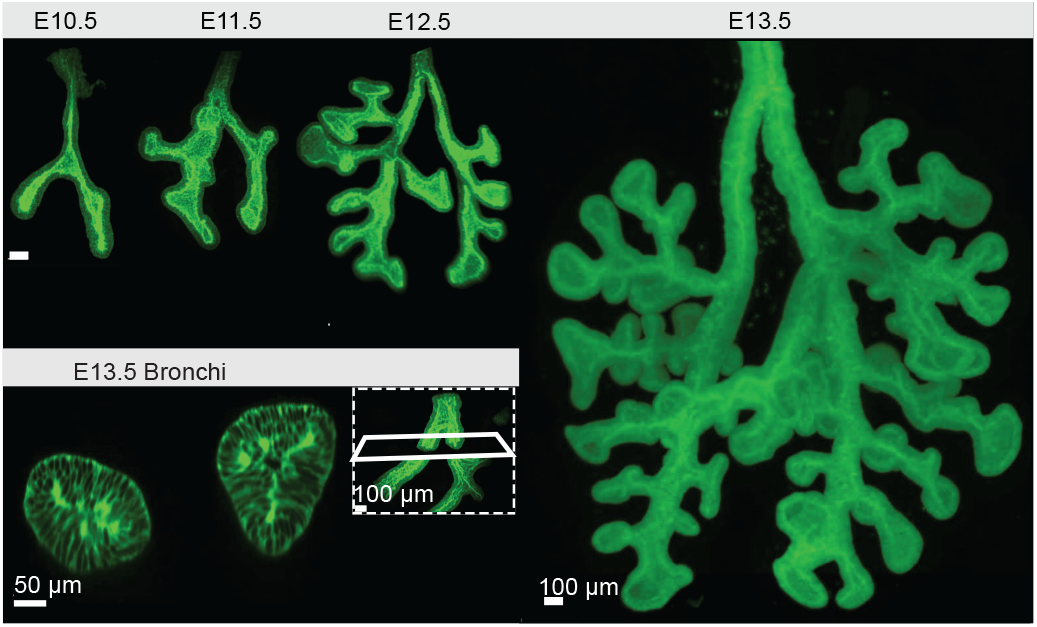
Developmental time course of mouse bronchial epithelial trees. The E10.5–E13.5 lungs are shown on the same scale, illustrating the strong enlargement between E12.5 and E13.5. Inset: Cross-section of E13.5 bronchi reveals a narrow lumen. Images reproduced from [30].

**Fig. S3:**
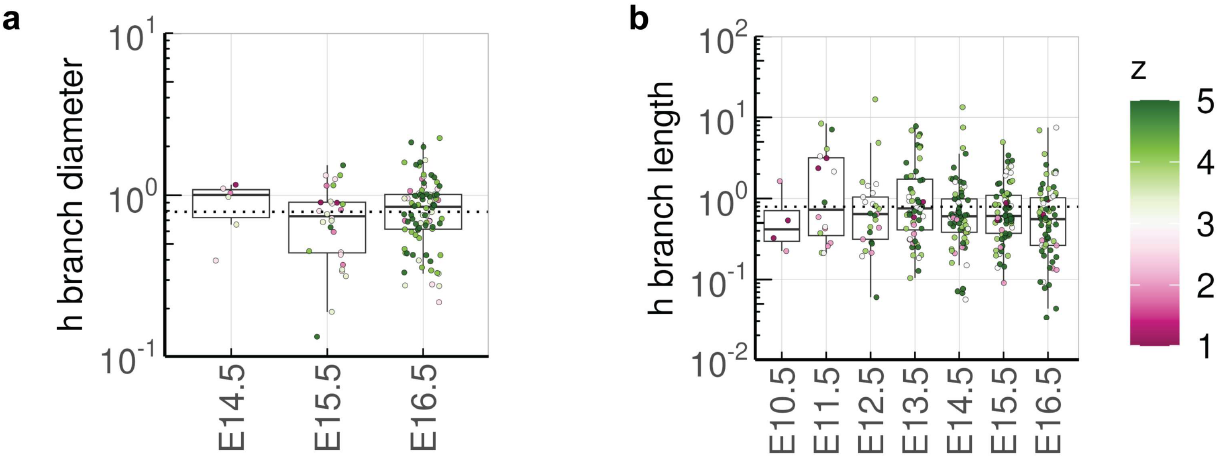
Morphometrics of mouse embryonic lungs. **a**,**b**, The distribution of homothety ratios of the inner branch diameters, *D*_*z*_ (**a**), and of the lengths, *L*_*z*_ (**b**), of all branches up to generation *z* = 6 over developmental time. *h* = 2^−1/3^ (dotted lines) for comparison. The data for E10.5–E13.5 was obtained using images published in [30].

**Fig. S4:**
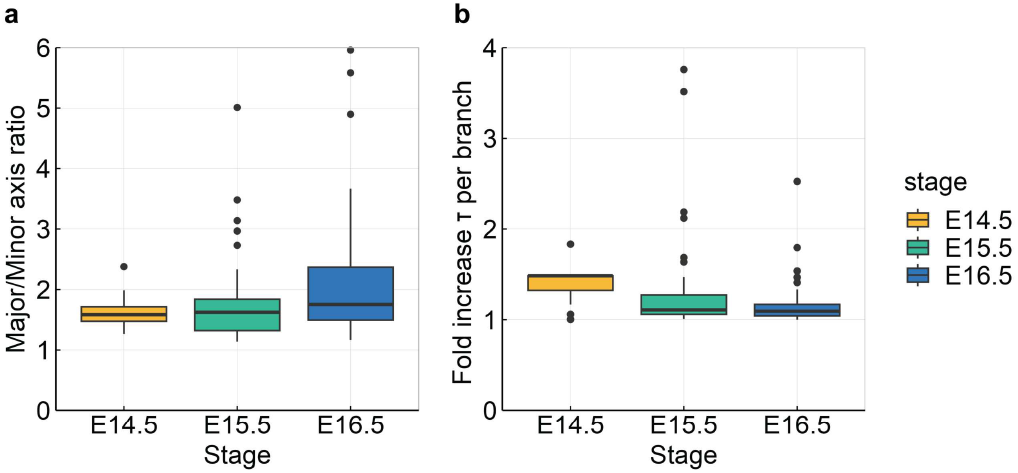
Impact of deviations from a cylindrical shape on wall shear stress. **a**, Circularity of tube cross-sections, quantified by the distribution of the major-minor axis length ratio of branch cross-sections. **b**, Impact of variable tube diameters along each branch on wall shear stress according to Eq. S2.

**Fig. S5:**
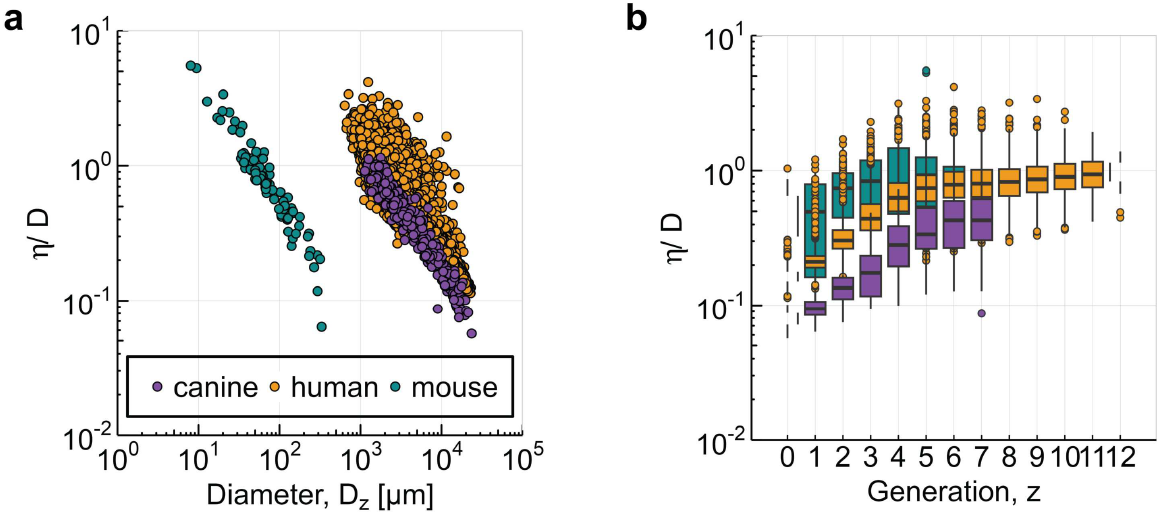
Lung branches can be approximated as thick cylinders. **a**, The ratio of wall thickness to inner diameter declines with increasing inner branch diameter. **b**, The ratio of wall thickness to inner diameter increases along the bronchial tree in all species.

**Fig. S6:**
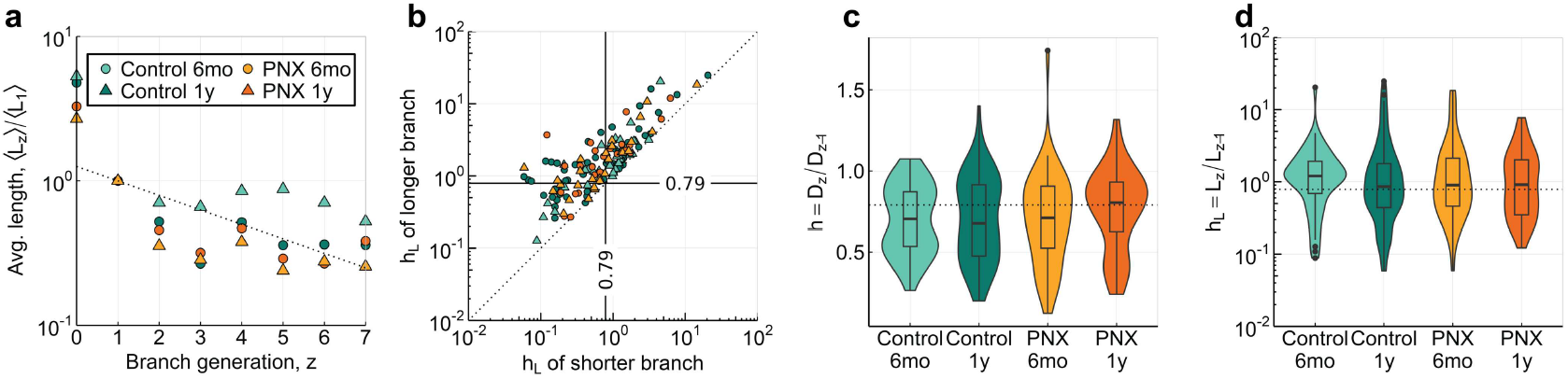
Morphometrics of canine lungs. **a**, Average branch lengths against branch generation. Dashed line represents the Hess–Murray law. **b**, Homothety ratio of the branch lengths of sister branches. **c**,**d** Variability of the homothety ratio of the inner branch diameters **(c)** and branch length **(d)**. *h* = 2^−1/3^ (dotted lines) for comparison. Data re-used from [53].

**Fig. S7:**
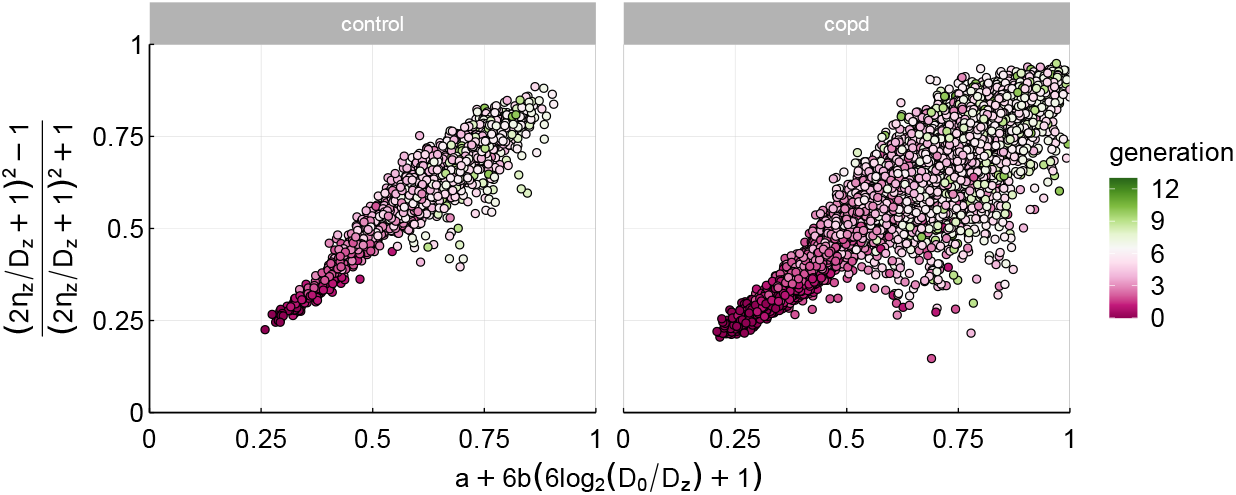
Extraction of the biophysical parameters in the human dataset.

**Fig. S8:**
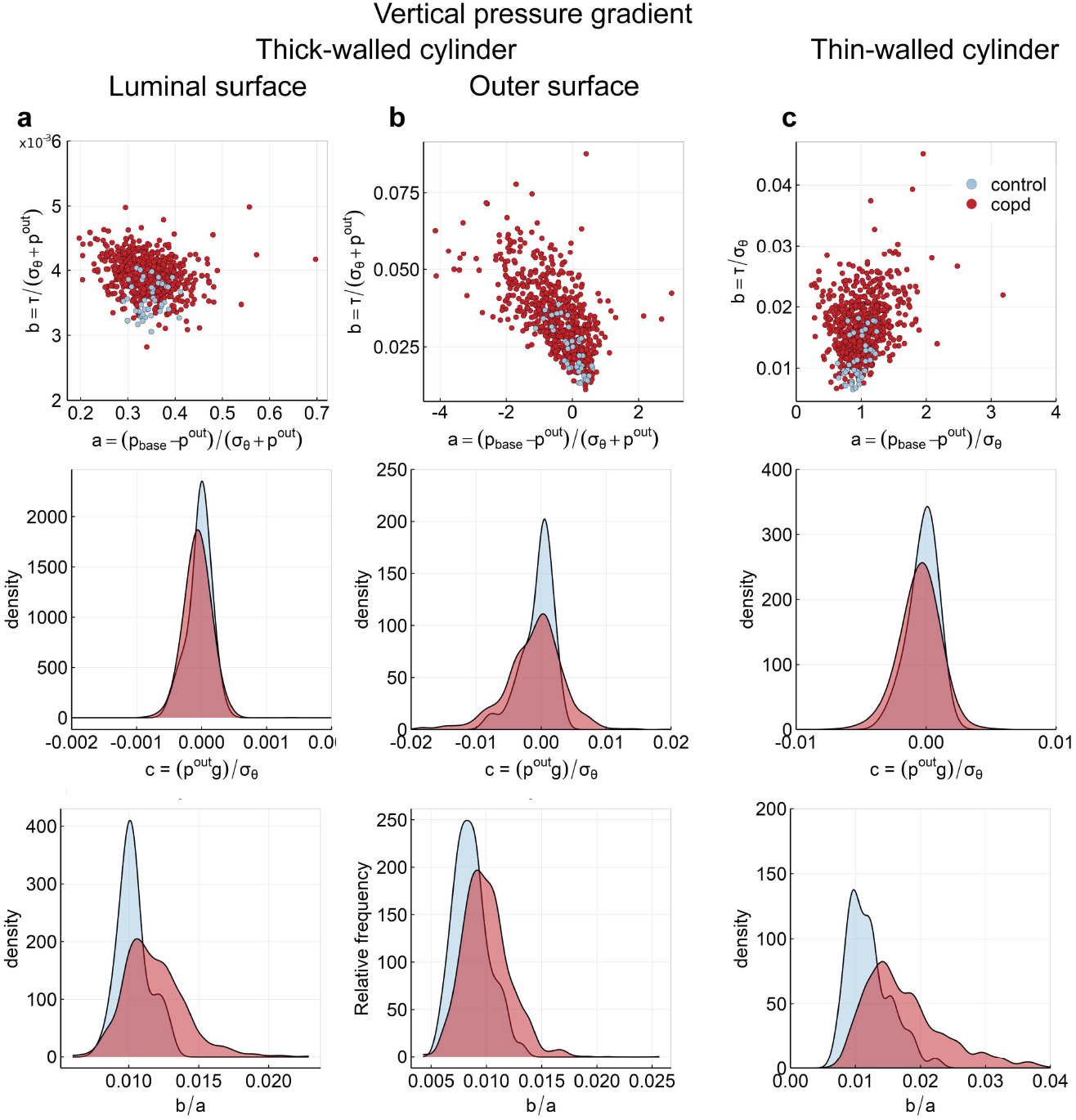
Distributions of inferred biophysical parameters for different models including a vertical pressure gradient. For a thick-walled cylinder on the luminal (**a**) (Eq. S25 and the outer wall surface (**b**) (Eq. S26 and a thin-walled cylinder (**c**).

**Fig. S9:**
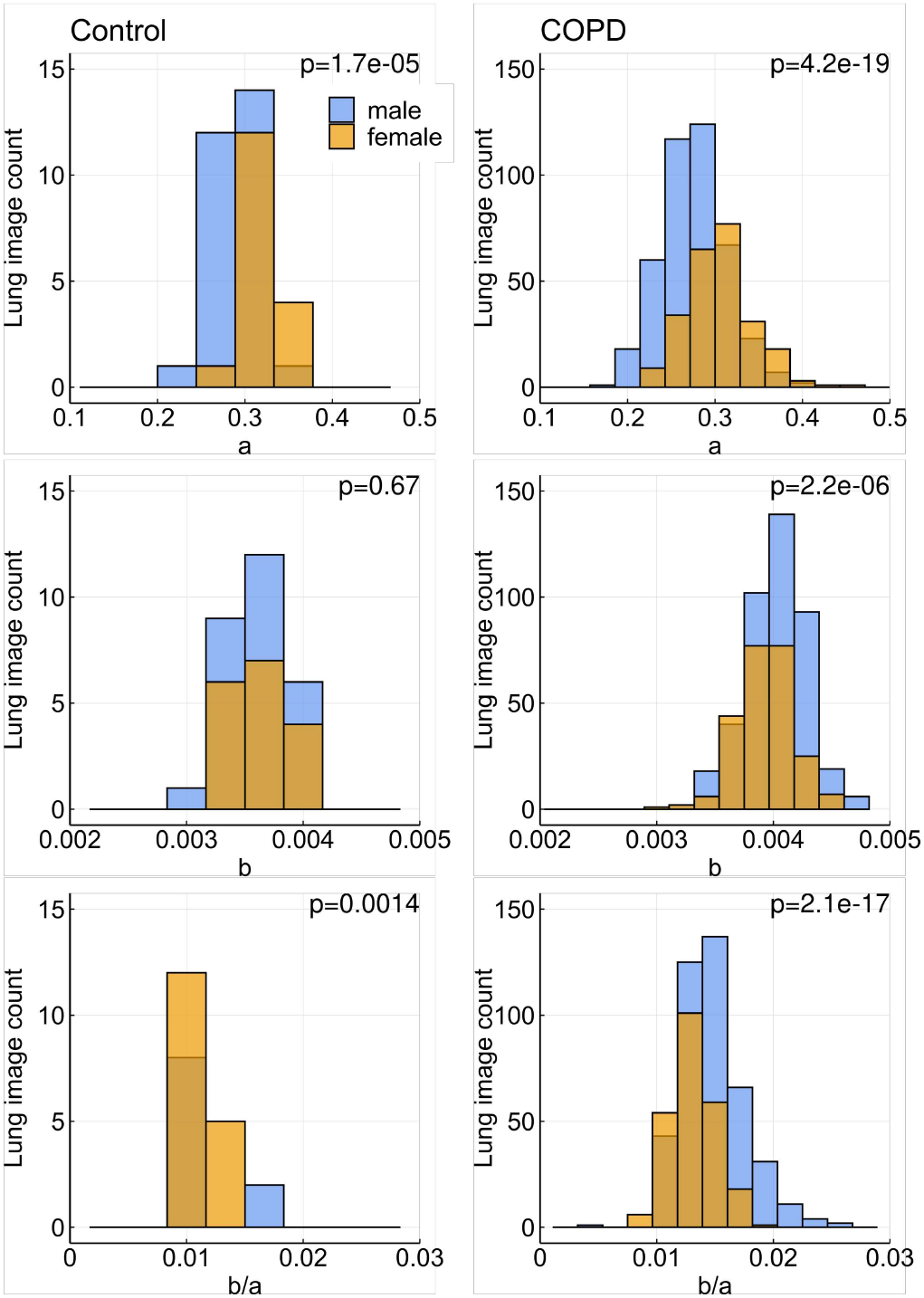
Dependency of biophysical parameters on patient sex. The empirical distributions of *a* = (*p*_base_ − *p*^out^)/*σ*_*θ*_ and *b*/*a* = *τ/*(*p*_base_ − *p*^out^) show statistically significant difference between male and female control patients (left, *n* = 45) and a significant difference between male and female COPD patients (right, *n* = 796). There is no significant difference in the *b* = *τ/σ*_*θ*_ distribution in control patients, but we find a significant difference in COPD patients. *p* values from Wilcoxon–Mann–Whitney tests.

## Supplementary Tables

**Table S1:**
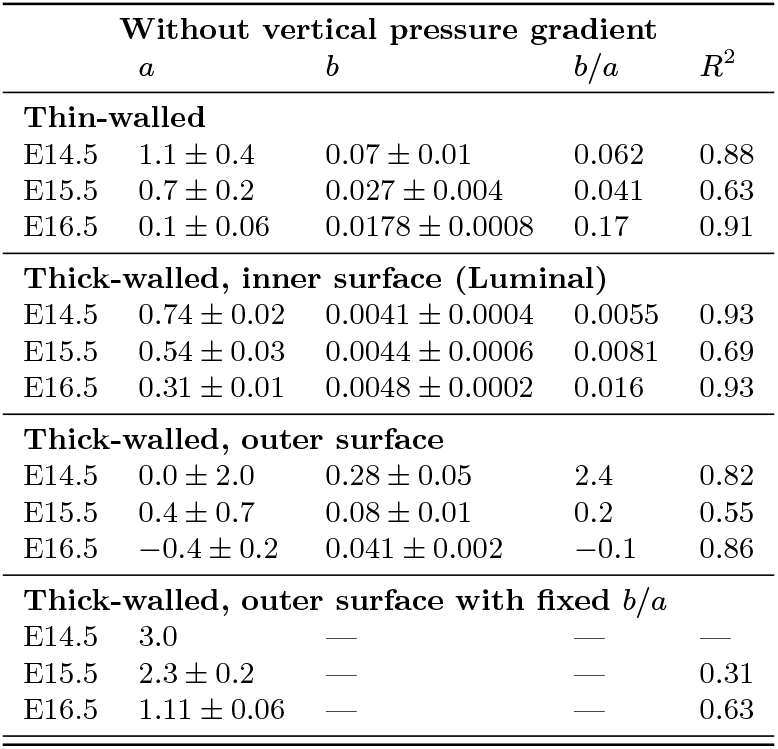
Biophysical parameters inferred for the developing mouse lung. Eq. S4 is fitted to the data shown in Fig. 5e. Uncertainties are s.e.

**Table S2:** Biophysical parameters inferred for the canine lung. Values are provided for different fit types and groups. Uncertainties are s.e.

| Without vertical pressure gradient |  |  |  |  |  |  |
| --- | --- | --- | --- | --- | --- | --- |
| | $a$ | $b$ | $c$ [1/mm] | $b/a$ | $c/a$ [1/mm] | $R^2$ |
| **Thin-walled** |  |  |  |  |  |  |
| Control 1year | $-0.1 \pm 0.04$ | $0.0113 \pm 0.0004$ | — | $-0.11$ | — | 0.83 |
| Control 6mo | $0.07 \pm 0.04$ | $0.0095 \pm 0.0006$ | — | 0.15 | — | 0.77 |
| Pnx 1year | $-0.1 \pm 0.04$ | $0.0093 \pm 0.0004$ | — | $-0.093$ | — | 0.9 |
| Pnx 6mo | $-0.25 \pm 0.07$ | $0.0138 \pm 0.0007$ | — | $-0.055$ | — | 0.86 |
| **Thick-walled, inner surface (Luminal)** |  |  |  |  |  |  |
| Control 1year | $0.12 \pm 0.01$ | $0.0044 \pm 0.0001$ | — | 0.036 | — | 0.92 |
| Control 6mo | $0.15 \pm 0.02$ | $0.0049 \pm 0.0003$ | — | 0.033 | — | 0.84 |
| Pnx 1year | $0.11 \pm 0.01$ | $0.0046 \pm 0.0002$ | — | 0.043 | — | 0.93 |
| Pnx 6mo | $0.18 \pm 0.01$ | $0.0048 \pm 0.0002$ | — | 0.026 | — | 0.94 |
| **Thick-walled, outer surface** |  |  |  |  |  |  |
| Control 1year | $-0.5 \pm 0.1$ | $0.022 \pm 0.001$ | — | $-0.043$ | — | 0.74 |
| Control 6mo | $-0.04 \pm 0.08$ | $0.015 \pm 0.001$ | — | $-0.35$ | — | 0.71 |
| Pnx 1year | $-0.32 \pm 0.07$ | $0.0154 \pm 0.0009$ | — | $-0.048$ | — | 0.85 |
| Pnx 6mo | $-0.9 \pm 0.2$ | $0.028 \pm 0.002$ | — | $-0.031$ | — | 0.79 |
| **Thick-walled, outer surface with fixed $b/a$** |  |  |  |  |  |  |
| Control 1year | $0.38 \pm 0.01$ | — | — | — | — | 0.59 |
| Control 6mo | $0.3 \pm 0.01$ | — | — | — | — | 0.62 |
| Pnx 1year | $0.127 \pm 0.005$ | — | — | — | — | 0.74 |
| Pnx 6mo | $0.37 \pm 0.02$ | — | — | — | — | 0.61 |
| With vertical pressure gradient |  |  |  |  |  |  |
| | $a$ | $b$ | $c$ [1/mm] | $b/a$ | $c/a$ [1/mm] | $R^2$ |
| **Thin-walled** |  |  |  |  |  |  |
| Control 1year | $-0.1 \pm 0.04$ | $0.0117 \pm 0.0005$ | $(1.1 \pm 6) \times 10^{-7}$ | $-0.11$ | $-1.1 \times 10^{-5}$ | 0.83 |
| Control 6mo | $0.07 \pm 0.05$ | $0.0096 \pm 0.0007$ | $(6 \pm 6) \times 10^{-7}$ | 0.13 | $8 \times 10^{-6}$ | 0.76 |
| Pnx 1year | $-0.1 \pm 0.04$ | $0.0094 \pm 0.0005$ | $(4 \pm 5) \times 10^{-7}$ | $-0.092$ | $-3.5 \times 10^{-6}$ | 0.9 |
| Pnx 6mo | $-0.28 \pm 0.07$ | $0.014 \pm 0.0007$ | $(-6 \pm 8) \times 10^{-7}$ | $-0.05$ | $2.1 \times 10^{-6}$ | 0.86 |
| **Thick-walled, inner surface** |  |  |  |  |  |  |
| Control 1year | $0.14 \pm 0.01$ | $0.0044 \pm 0.0001$ | $(1 \pm 2) \times 10^{-7}$ | 0.032 | $6.2 \times 10^{-7}$ | 0.91 |
| Control 6mo | $0.14 \pm 0.02$ | $0.0049 \pm 0.0003$ | $(1 \pm 2) \times 10^{-7}$ | 0.034 | $9.6 \times 10^{-7}$ | 0.83 |
| Pnx 1year | $0.05 \pm 0.02$ | $0.0047 \pm 0.0002$ | $(1 \pm 2 \times 10^{-7})$ | 0.085 | $1.2 \times 10^{-6}$ | 0.93 |
| Pnx 6mo | $0.12 \pm 0.01$ | $0.0048 \pm 0.0002$ | $(-3 \pm 2) \times 10^{-7}$ | 0.041 | $-2.2 \times 10^{-6}$ | 0.94 |
| **Thick-walled, outer surface** |  |  |  |  |  |  |
| Control 1year | $-0.5 \pm 0.1$ | $0.023 \pm 0.001$ | $(3 \pm 1) \times 10^{-6}$ | $-0.044$ | $-6.2 \times 10^{-6}$ | 0.75 |
| Control 6mo | $-0.02 \pm 0.09$ | $0.016 \pm 0.001$ | $(1 \pm 1) \times 10^{-6}$ | $-0.64$ | $-5.6 \times 10^{-5}$ | 0.7 |
| Pnx 1year | $-0.33 \pm 0.08$ | $0.0157 \pm 0.0009$ | $(1 \pm 1) \times 10^{-6}$ | $-0.048$ | $-2.2 \times 10^{-6}$ | 0.85 |
| Pnx 6mo | $-1.0 \pm 0.2$ | $0.029 \pm 0.002$ | $(-1 \pm 2) \times 10^{-6}$ | $-0.029$ | $1.2 \times 10^{-6}$ | 0.79 |
| **Thick-walled, outer surface with fixed $b/a$** |  |  |  |  |  |  |
| Control 1year | $0.64 \pm 0.02$ | — | $(7 \pm 2) \times 10^{-6}$ | — | $1.1 \times 10^{-5}$ | 0.74 |
| Control 6mo | $0.51 \pm 0.02$ | — | $(2 \pm 2) \times 10^{-6}$ | — | $4 \times 10^{-6}$ | 0.68 |
| Pnx 1year | $0.195 \pm 0.008$ | — | $(-1 \pm 5) \times 10^{-6}$ | — | $-5.1 \times 10^{-6}$ | 0.86 |
| Pnx 6mo | $0.61 \pm 0.02$ | — | $(-4 \pm 4) \times 10^{-6}$ | — | $-6.6 \times 10^{-6}$ | 0.78 |

**Table S3:** Biophysical parameters inferred for the adults human lung. Eq. S4 is fitted to each patient. Shown are the mean parameters and fit qualities for COPD patients and controls. Uncertainties are standard deviations of the mean parameters

| **Without vertical pressure gradient** |  |  |  |  |  |  |
| --- | --- | --- | --- | --- | --- | --- |
| | $a$ | $b$ | $c$ [1/mm] | $b/a$ | $c/a$ [1/mm] | $R^2$ |
| **Thin-walled** |  |  |  |  |  |  |
| control | $0.1 \pm 0.1$ | $(1.0 \pm 0.3) \times 10^{-2}$ | --- | $-0.04 \pm 0.79$ | --- | 0.86 |
| copd | $-0.01 \pm 0.181$ | $(1.5 \pm 0.5) \times 10^{-2}$ | --- | $0.3 \pm 11.0$ | --- | 0.85 |
| **Thick-walled, inner surface (Luminal)** |  |  |  |  |  |  |
| control | $0.27 \pm 0.05$ | $(3.6 \pm 0.2) \times 10^{-3}$ | --- | $0.014 \pm 0.003$ | --- | 0.91 |
| copd | $0.29 \pm 0.05$ | $(4.0 \pm 0.4) \times 10^{-3}$ | --- | $0.01 \pm 0.17$ | --- | 0.92 |
| **Thick-walled, outer surface** |  |  |  |  |  |  |
| control | $-0.1 \pm 0.3$ | $(2.0 \pm 0.8) \times 10^{-2}$ | --- | $0.01 \pm 0.42$ | --- | 0.82 |
| copd | $-0.7 \pm 0.6$ | $(3.0 \pm 1.0) \times 10^{-2}$ | --- | $-0.09 \pm 2.55$ | --- | 0.77 |
| **Thick-walled, outer surface with fixed $b/a$** |  |  |  |  |  |  |
| control | $0.8 \pm 0.3$ | --- | --- | --- | --- | 0.65 |
| copd | $1.2 \pm 0.4$ | --- | --- | --- | --- | 0.56 |
| **Thin-walled with vertical pressure gradient** |  |  |  |  |  |  |
| control | $0.2 \pm 0.1$ | $(1.0 \pm 0.3) \times 10^{-2}$ | $(-1 \pm 8) \times 10^{-5}$ | $0.04 \pm 0.27$ | $(-9 \pm 207) \times 10^{-5}$ | 0.86 |
| copd | $0.1 \pm 0.3$ | $(1.5 \pm 0.3) \times 10^{-2}$ | $(-3 \pm 15) \times 10^{-5}$ | $-0.02 \pm 1.49$ | $(9 \pm 103) \times 10^{-4}$ | 0.85 |
| **Thick-walled, inner surface with vertical pressure gradient** |  |  |  |  |  |  |
| control | $0.31 \pm 0.05$ | $(3.6 \pm 0.2) \times 10^{-3}$ | $(-2 \pm 17) \times 10^{-6}$ | $0.012 \pm 0.002$ | $(-1 \pm 60) \times 10^{-5}$ | 0.9 |
| copd | $0.34 \pm 0.05$ | $(3.9 \pm 0.3) \times 10^{-3}$ | $(-5 \pm 22) \times 10^{-6}$ | $0.012 \pm 0.002$ | $(-2 \pm 70) \times 10^{-5}$ | 0.92 |
| **Thick-walled, outer surface with vertical pressure gradient** |  |  |  |  |  |  |
| control | $0.02 \pm 0.36$ | $(2.0 \pm 0.8) \times 10^{-2}$ | $(-4 \pm 20) \times 10^{-4}$ | $-0.02 \pm 0.58$ | $(3 \pm 17) \times 10^{-3}$ | 0.82 |
| copd | $-0.4 \pm 0.8$ | $(3.0 \pm 1.0) \times 10^{-2}$ | $(-8 \pm 46) \times 10^{-4}$ | $-0.02 \pm 1.02$ | $(8 \pm 69) \times 10^{-3}$ | 0.78 |
| **Thick-walled, outer surface with fixed $b/a$ and vertical pressure gradient** |  |  |  |  |  |  |
| control | $1.6 \pm 0.7$ | --- | $(-4 \pm 8) \times 10^{-4}$ | --- | $(-3 \pm 6) \times 10^{-4}$ | 0.81 |
| copd | $2.4 \pm 0.9$ | --- | $(4 \pm 11) \times 10^{-4}$ | --- | $(1 \pm 6) \times 10^{-4}$ | 0.76 |

## References

[1] R. R. Beichel, R. W. Glenny, C. Bauer, and M. A. Krueger. Lung Anatomy + Particle Deposition (lapd) Mouse Archive. https://iro.uiowa.edu/esploro/outputs/dataset/9983559095702771, 2019.

[2] R. W. Glenny, M. Krueger, C. Bauer, and R. R. Beichel. The fractal geometry of bronchial trees differs by strain in mice. J. Appl. Physiol., 128:362–367, 2020. doi: 10.1152/japplphysiol.00838.2019.

[3] A. W. Sheel, J. A. Guenette, R. Yuan, L. Holy, J. R. Mayo, A. M. McWilliams, S. Lam, and H. O. Coxson. Evidence for dysanapsis using computed tomographic imaging of the airways in older ex-smokers. J. Appl. Physiol., 107:1622–1628, 2009. doi: 10.1152/japplphysiol.00562.2009.

[4] E. R. Weibel and D.M. Gomez. Architecture of the human lung. Use of quantitative methods establishes fundamental relations between size and number of lung structures. Science, 137:577–585, 1962. doi: 10.1126/science.137.3530.577.

[5] K. Horsfield and G. Cumming. Morphology of the bronchial tree in man. J. Appl. Physiol., 24:373–383, 1968. doi: 10.1152/jappl.1968.24.3.373.

[6] W. R. Hess. Das Prinzip des kleinsten Kraftverbrauchs im Dienste hämodynamischer Forschung. Archiv Anat. Physiol., pages 1–62, 1914.

[7] C. D. Murray. The Physiological Principle of Minimum Work: I. The Vascular System and the Cost of Blood Volume. Proc. Natl. Acad. Sci. U. S. A., 12:207–214, 1926. doi: 10.1073/pnas.12.3.207.

[8] T. Wilson. Design of the Bronchial Tree. Nature, 213:668–669, 1967. doi: 10.1038/213668a0.

[9] K. K. Bokka, E. C. Jesudason, O. A. Lozoya, F. Guilak, D. Warburton, and S. R. Lubkin. Morphogenetic Implications of Peristalsis-Driven Fluid Flow in the Embryonic Lung. PLOS ONE, 10:e0132015, 2015. doi: 10.1371/journal.pone.0132015.

[10] A. Bejan, L. A. O. Rocha, and S. Lorente. Thermodynamic optimization of geometry: T-and Y-shaped constructs of fluid streams. Int. J. Therm. Sci., 39:949–960, 2000. doi: 10.1016/S1290-0729(00)01176-5.

[11] E. R. Weibel. Morphometry of the Human Lung. Springer Berlin, Heidelberg, 1963. ISBN 978-3-642-87553-3. doi: 10.1007/978-3-642-87553-3.

[12] K. Horsfield and G. Cumming. Angles of branching and diameters of branches in the human bronchial tree. Bull. Math. Biophys., 29:245–259, 1967. doi: 10.1007/BF02476898.

[13] O. G. Raabe, H. C. Yeah, G. M. Schum, and R. F. Phalen. Trachobronchial geometry: human, dog, rat, hamster. Technical report, Lovelace Foundation for Medical Education and Research, 1976. Report LF-53.

[14] R. F. Phalen, H. C. Yeh, G. M. Schum, and O. G. Raabe. Application of an idealized model to morphometry of the mammalian tracheobronchial tree. Anat. Rec, 190:167–176, 1978. doi: 10.1002/ar.1091900202.

[15] H. C. Yeh, G. M. Schum, and M. T. Duggan. Anatomic models of the tracheobronchial and pulmonary regions of the rat. Anat. Rec., 195:483–492, 1979. doi: 10.1002/ar.1091950308.

[16] A. Thurlbeck and K. Horsfield. Branching angles in the bronchial tree related to order of branching. Respir. Physiol., 41:173–181, 1980. doi: 10.1016/0034-5687(80)90050-X.

[17] M. H. Tawhai, P. Hunter, J. Tschirren, J. Reinhardt, G. McLennan, and E. A. Hoffman. CT-based geometry analysis and finite element models of the human and ovine bronchial tree. J. Appl. Physiol., 97:2310–2321, 2004. doi: 10.1152/japplphysiol.00520.2004.

[18] W. B. Counter, I. Q. Wang, T. H. Farncombe, and N. R. Labiris. Airway and pulmonary vascular measurements using contrast-enhanced micro-CT in rodents. Am. J. Physiol. - Lung Cell. Mol. Physiol., 304:L831–L843, 2013. doi: 10.1152/ajplung.00281.2012.

[19] D. Hunt and V. M. Savage. Asymmetries arising from the space-filling nature of vascular networks. Phys. Rev. E, 93:062305, 2016. doi: 10.1103/PhysRevE.93.062305.

[20] T. Van de Moortele, C. H. Wendt, and F. Coletti. Morphological and functional properties of the conducting human airways investigated by in vivo computed tomography and in vitro MRI. J. Appl. Physiol., 124:400–413, 2018. doi: 10.1152/japplphysiol.00490.2017.

[21] K. Onuma, M. Ebina, T. Takahashi, and T. Nukiwa. Irregularity of airway branching in a mouse bronchial tree: a 3-D morphometric study. Tohoku J. Exp. Med., 194:157–164, 2001. doi: 10.1620/tjem.194.157.

[22] B. J. West, V. Bhargava, and A. L. Goldberger. Beyond the principle of similitude: renormalization in the bronchial tree. J. Appl. Physiol., 60:1089–1097, 1986.

[23] M. Florens, B. Sapoval, and M. Filoche. Optimal branching asymmetry of hydrodynamic pulsatile trees. Phys. Rev. Lett., 106:178104, 2011. doi: 10.1103/PhysRevLett.106.178104.

[24] A. Majumdar, A. M. Alencar, S. V. Buldyrev, Z. Hantos, K. R. Lutchen, H. E. Stanley, and B. Suki. Relating airway diameter distributions to regular branching asymmetry in the lung. Phys. Rev. Lett., 95:168101, 2005. doi: 10.1103/PhysRevLett.95.168101.

[25] H. B. Uylings. Optimization of diameters and bifurcation angles in lung and vascular tree structures. Bull. Math. Biol., 39:509–520, 1977. doi: 10.1007/BF02461198.

[26] D. Iber. The control of lung branching morphogenesis. Curr. Top. Dev. Biol., 143:205–237, 2021. doi: 10.1016/bs.ctdb.2021.02.002.

[27] R. J. Metzger, O. D. Klein, G. R. Martin, and M. A. Krasnow. The branching programme of mouse lung development. Nature, 453:745–750, 2008. doi: 10.1038/nature07005.

[28] E. E. Morrisey and B. L. Hogan. Preparing for the first breath: genetic and cellular mechanisms in lung development. Dev. Cell, 18:8–23, 2010. doi: 10.1016/j.devcel.2009.12.010.

[29] S. F. Barre, D. Haberthur, T. P. Cremona, M. Stampanoni, and J. C. Schittny. The total number of acini remains constant throughout postnatal rat lung development. Am. J. Physiol. - Lung Cell. Mol. Physiol., 311:L1082–L1089, 2016. doi: 10.1152/ajplung.00325.2016.

[30] L. Conrad, S. V. M. Runser, H. Fernando Gómez, C. M. Lang, M. S. Dumond, A. Sapala, L. Schaumann, O. Michos, R. Vetter, and D. Iber. The biomechanical basis of biased epithelial tube elongation in lung and kidney development. Development, 148:dev194209, 2021. doi: 10.1242/dev.194209.

[31] K. Kishimoto, M. Tamura, M. Nishita, Y. Minami, A. Yamaoka, T. Abe, M. Shigeta, and M. Morimoto. Synchronized mesenchymal cell polarization and differentiation shape the formation of the murine trachea and esophagus. Nat. Commun., 9:2816, 2018. doi: 10.1038/s41467-018-05189-2.

[32] U. Pohl, J. Holtz, R. Busse, and E. Bassenge. Crucial role of endothelium in the vasodilator response to increased flow in vivo. Hypertens., 8:37–44, 1986. doi: 10.1161/01.hyp.8.1.37.

[33] J. R. Hove, R. W. Köster, A. S. Forouhar, G. Acevedo-Bolton, S. E. Fraser, and M. Gharib. Intracardiac fluid forces are an essential epigenetic factor for embryonic cardiogenesis. Nature, 421:172–177, 2003. doi: 10.1038/nature01282.

[34] S. Marbach, N. Ziethen, L. Bastin, F. K. Bäuerle, and K. Alim. Vein fate determined by flow-based but time-delayed integration of network architecture. eLife, 12:e78100, 2023. doi: 10.7554/eLife.78100.

[35] S. Marbach, N. Ziethen, and K. Alim. Vascular adaptation model from force balance: Physarum polycephalum as a case study. New J. Phys., 25:123052, 2023. doi: 10.1088/1367-2630/ad1488.

[36] G. N. Karam. Biomechanical Model of the Xylem Vessels in Vascular Plants. Ann. Bot., 95:1179–1186, 2005. doi: 10.1093/aob/mci130.

[37] C. M. Nelson, J. P. Gleghorn, M.-F. Pang, J. M. Jaslove, K. Goodwin, V. D. Varner, E. Miller, D. C. Radisky, and H. A. Stone. Microfluidic chest cavities reveal that transmural pressure controls the rate of lung development. Development, 144:4328–4335, 2017. doi: 10.1242/dev.154823.

[38] R. Harding and S. B. Hooper. Regulation of lung expansion and lung growth before birth. J. Appl. Physiol., 81:209–224, 1996. doi: 10.1152/jappl.1996.81.1.209.

[39] J. H. Cartwright, O. Piro, and I. Tuval. Fluid dynamics in developmental biology: moving fluids that shape ontogeny. HFSP J., 3:77–93, 2009. doi: 10.2976/1.3043738.

[40] R. Harding, A. D. Bocking, and J. N. Sigger. Influence of upper respiratory tract on liquid flow to and from fetal lungs. J. Appl. Physiol., 61:68–74, 1986. doi: 10.1152/jappl.1986.61.1.68.

[41] R. Harding, S. B. Hooper, and V. K. M. Han. Abolition of fetal breathing movements by spinal cord transection leads to reductions in fetal lung liquid volume, lung growth, and IGF-II gene expression. Pediatr. Res., 34:148–153, 1993. doi: 10.1203/00006450-199308000-00008.

[42] M. Zamir. Shear forces and blood vessel radii in the cardiovascular system. J. Gen. Physiol., 69:449–461, 1977. doi: 10.1085/jgp.69.4.449.

[43] R. Tarran, B. Button, M. Picher, A. M. Paradiso, C. M. Ribeiro, E. R. Lazarowski, L. Zhang, P. L. Collins, R. J. Pickles, J. J. Fredberg, and R. C. Boucher. Normal and cystic fibrosis airway surface liquid homeostasis. The effects of phasic shear stress and viral infections. J. Biol. Chem., 280:35751–35759, 2005. doi: 10.1074/jbc.M505832200.

[44] K. Short, M. Hodson, and I. Smyth. Spatial mapping and quantification of developmental branching morphogenesis. Development, 140:471–478, 2013. doi: 10.1242/dev.088500.

[45] S. Fujii, T. Muranaka, J. Matsubayashi, S. Yamada, A. Yoneyama, and T. Takakuwa. Bronchial tree of the human embryo: Categorization of the branching mode as monopodial and dipodial. PLOS ONE, 16:e0245558, 2021. doi: 10.1371/journal.pone.0245558.

[46] S. Fujii, T. Muranaka, J. Matsubayashi, S. Yamada, A. Yoneyama, and T. Takakuwa. Bronchial tree of the human embryo: Examination based on a mammalian model. J. Anat., 244:159–169, 2024. doi: 10.1111/joa.13946.

[47] S. J. Lai-Fook and J. R. Rodarte. Pleural pressure distribution and its relationship to lung volume and interstitial pressure. J. Appl. Physiol., 70:967–978, 1991. doi: 10.1152/jappl.1991.70.3.967.

[48] S. J. Lai-Fook. Pleural mechanics and fluid exchange. Physiol. Rev., 84:385–410, 2004. doi: 10.1152/physrev.00026.2003.

[49] A. Yartsev. Vertical gradient of pleural pressure, 2019. URL https://derangedphysiology.com/main/cicm-primary-exam/required-reading/respiratory-system/Chapter%200356/vertical-gradient-pleural-pressure.

[50] R. E. Olver, E. E. Schneeberger, and D. V. Walters. Epithelial solute permeability, ion transport and tight junction morphology in the developing lung of the fetal lamb. J. Physiol., 315:395–412, 1981. doi: 10.1113/jphysiol.1981.sp013754.

[51] S. B. Hooper and R. Harding. Fetal lung liquid: a major determinant of the growth and functional development of the fetal lung. Clin. Exp. Pharmacol. Physiol., 22:235–241, 1995. doi: 10.1111/j.1440-1681.1995.tb01988.x.

[52] D. E. Olson, G. A. Dart, and G. F. Filley. Pressure drop and fluid flow regime of air inspired into the human lung. J. Appl. Physiol., 28:482–494, 1970. doi: 10.1152/jappl.1970.28.4.482.

[53] D. M. Dane, Jr. Johnson, R. L., and C. C. Hsia. Dysanaptic growth of conducting airways after pneumonectomy assessed by CT scan. J. Appl. Physiol., 93:1235–1242, 2002. doi: 10.1152/japplphysiol.00970.2001.

[54] K. Tsujikawa, R. Muramatsu, and T. Miyata. CSF pressure in fetal mice in utero: External factors pressurize the intraventricular space. bioRxiv, 2024. doi: 10.1101/2024.09.08.611845.

[55] P. Ravikumar, C. Yilmaz, D. M. Dane, D. J. Bellotto, A. S. Estrera, and C. C. W. Hsia. Defining a stimuli-response relationship in compensatory lung growth following major resection. J. Appl. Physiol., 116:816–824, 2014. doi: 10.1152/japplphysiol.01291.2013.

[56] B. B. Ross. Influence of bronchial tree structure on ventilation in the dog’s lung as inferred from measurements of a plastic cast. J. Appl. Physiol., 10:1–14, 1957. doi: 10.1152/jappl.1957.10.1.1.

[57] T. Weikert, L. Friebe, A. Wilder-Smith, S. Yang, J. I. Sperl, D. Neumann, A. Balachandran, J. Bremerich, and A. W. Sauter. Automated quantification of airway wall thickness on chest CT using retina U-Nets - Performance evaluation and application to a large cohort of chest CTs of COPD patients. Eur. J. Radiol., 155:110460, 2022. doi: 10.1016/j.ejrad.2022.110460.

[58] S. Chen, M. Kuhn, K. Prettner, F. Yu, T. Yang, T. Barnighausen, D. E. Bloom, and C. Wang. The global economic burden of chronic obstructive pulmonary disease for 204 countries and territories in 2020-50: a health-augmented macroeconomic modelling study. Lancet Glob. Health, 11:e1183–e1193, 2023. doi: 10.1016/S2214-109X(23)00217-6.

[59] A. Agusti, B. R. Celli, G. J. Criner, D. Halpin, A. Anzueto, P. Barnes, J. Bourbeau, M. K. Han, F. J. Martinez, M. Montes de Oca, K. Mortimer, A. Papi, I. Pavord, N. Roche, S. Salvi, D. D. Sin, D. Singh, R. Stockley, M. V. Lopez Varela, J. A. Wedzicha, and C. F. Vogelmeier. Global Initiative for Chronic Obstructive Lung Disease 2023 Report: GOLD Executive Summary. Am. J. Respir. Crit. Care Med., 207:819–837, 2023. doi: 10.1164/rccm.202301-0106PP.

[60] M. Montaudon, P. Desbarats, P. Berger, G. de Dietrich, R. Marthan, and F. Laurent. Assessment of bronchial wall thickness and lumen diameter in human adults using multi-detector computed tomography: comparison with theoretical models. J. Anat., 211:579–588, 2007. doi: 10.1111/j.1469-7580.2007.00811.x.

[61] J. B. West and P. Hugh-Jones. Patterns of gas flow in the upper bronchial tree. J. Appl. Physiol., 14:753–759, 1959. doi: 10.1152/jappl.1959.14.5.753.

[62] T. J. Pedley, R. C. Schroter, and M. F. Sudlow. The prediction of pressure drop and variation of resistance within the human bronchial airways. Respir. Physiol., 9:387–405, 1970. doi: 10.1016/0034-5687(70)90094-0.

[63] J. P. O’Connor, B. M. Hanley, T. I. Mulcahey, E. E. Sheets, and K. W. Shuey. N(2) gas egress from patients’ airways during LN(2) spray cryotherapy. Med. Eng. Phys., 44:63–72, 2017. doi: 10.1016/j.medengphy.2017.02.017.

[64] A. I. Nikiforov and R. B. Schlesinger. Morphometric variability of the human upper bronchial tree. Respir. Physiol., 59:289–299, 1985. doi: 10.1016/0034-5687(85)90134-3.

[65] P. J. Basser, T. A. McMahon, and P. Griffith. The Mechanism of Mucus Clearance in Cough. J. Biomech. Eng., 111:288–297, 1989. doi: 10.1115/1.3168381.

[66] B. Yeganeh, C. Bilodeau, and M. Post. Explant Culture for Studying Lung Development. In P. Delgado-Olguin, editor, Mouse Embryogenesis: Methods and Protocols, volume 1752, chapter 8, pages 81–90. Humana Press, 2018. doi: 10.1007/978-1-4939-7714-78.

[67] D. Menshykau, P. Blanc, E. Unal, V. Sapin, and D. Iber. An interplay of geometry and signaling enables robust lung branching morphogenesis. Development, 141:4526–4536, 2014. doi: 10.1242/dev.116202.

[68] B. P. Flannery, H. W. Deckman, W. G. Roberge, and K.L. D’Amico. Three-dimensional X-ray microtomography. Science, 237:1439–1444, 1987. doi: 10.1126/science.237.4821.1439.

[69] D. P. Clark and C. T. Badea. Micro-CT of rodents: State-of-the-art and future perspectives. Phys. Med., 30:619–634, 2014. doi: 10.1016/j.ejmp.2014.05.011.

[70] C. Aeby. Der Bronchialbaum der Säugethiere und des Menschen, nebst Bemerkungen über den Bronchialbaum der Vögel und Reptilien. Engelmann, Leipzig, 1880.

[71] M. Zheng, N. A. Borkar, Y. Yao, X. Ye, E. R. Vogel, C. M. Pabelick, and Y. S. Prakash. Mechanosensitive channels in lung disease. Front Physiol, 14:1302631, 2023. doi: 10.3389/fphys.2023.1302631.

[72] J. Yin and W. M. Kuebler. Mechanotransduction by TRP channels: general concepts and specific role in the vasculature. Cell Biochem. Biophys., 56:1–18, 2010. doi: 10.1007/s12013-009-9067-2.

[73] M. Thiriet. Mechanotransduction and vascular resistance. In P. Lanzer, editor, PanVascular Medicine. Springer, 2014. doi: 10.1007/978-3-642-37393-0258-2.

[74] B. J. McHugh, A. Murdoch, C. Haslett, and T. Sethi. Loss of the integrin-activating transmembrane protein Fam38A (Piezo1) promotes a switch to a reduced integrin-dependent mode of cell migration. PloS ONE, 7:e40346, 2012. doi: 10.1371/journal.pone.0040346.

[75] S. A. Gudipaty, J. Lindblom, P. D. Loftus, M. J. Redd, K. Edes, C. F. Davey, V. Krishnegowda, and J. Rosenblatt. Mechanical stretch triggers rapid epithelial cell division through Piezo1. Nature, 543:118–121, 2017. doi: 10.1038/nature21407.

[76] T. F. Sherman. On connecting large vessels to small. The meaning of Murray’s law. J. Gen. Physiol., 78:431–453, 1981. doi: 10.1085/jgp.78.4.431.

[77] S. R. Schachat, C. K. Boyce, J. L. Payne, and D. Lentink. Lepidoptera demonstrate the relevance of Murray’s Law to circulatory systems with tidal flow. BMC Biol., 19:204, 2021. doi: 10.1186/s12915-021-01130-0.

[78] D. Akita, I. Kunita, M. D. Fricker, S. Kuroda, K. Sato, and T. Nakagaki. Experimental models for Murray’s law. J. Phys. D: Appl. Phys., 50:024001, 2016. doi: 10.1088/1361-6463/50/2/024001.

[79] K. A. McCulloh, J. S. Sperry, and F. R. Adler. Water transport in plants obeys Murray’s law. Nature, 421:939–942, 2003. doi: 10.1038/nature01444.

[80] A. R. Pries, T. W. Secomb, and P. Gaehtgens. Design principles of vascular beds. Circ Res, 77(5):1017–23, 1995. doi: 10.1161/01.res.77.5.1017.

[81] R. S. Reneman and A. P. G. Hoeks. Wall shear stress as measured in vivo: consequences for the design of the arterial system. Med. Biol. Eng. Comput., 46:499–507, 2008. doi: 10.1007/s11517-008-0330-2.

[82] B. Mauroy and B. Moreau. Murray’s law revisited: Quémada’s fluids and fractal trees. J. Rheol., 59:1419–1430, 2015. doi: 10.1122/1.4934240.

[83] A. R. Pries, B. Reglin, and T. W. Secomb. Structural adaptation of vascular networks: role of the pressure response. Hypertension, 38(6):1476–9, 2001. doi: 10.1161/hy1201.100592.

[84] A. R. Pries and T. W. Secomb. Making microvascular networks work: angiogenesis, remodeling, and pruning. Physiology, 29:446–455, 2014. doi: 10.1152/physiol.00012.2014.

[85] L. A. Taber. Towards a unified theory for morphomechanics. Philos Trans A Math Phys Eng Sci, 367(1902):3555–83, 2009. doi: 10.1098/rsta.2009.0100. URL https://www.ncbi.nlm.nih.gov/pubmed/19657011 https://www.ncbi.nlm.nih.gov/pmc/articles/PMC2865877/pdf/rsta20090100.pdf.

[86] G. Genikhovich, P. Fried, M. M. Prunster, J. B. Schinko, A. F. Gilles, D. Fredman, K. Meier, D. Iber, and U. Technau. Axis patterning by bmps: Cnidarian network reveals evolutionary constraints. Cell Rep, 10(10):1646–1654, 2015.

[87] E. Hannezo, Clgj Scheele, M. Moad, N. Drogo, R. Heer, R. V. Sampogna, J. van Rheenen, and B. D. Simons. A unifying theory of branching morphogenesis. Cell, 171(1):242–255 e27, 2017.

[88] E. Gallo, S. De Renzis, J. Sharpe, R. Mayor, and J. Hartmann. Versatile system cores as a conceptual basis for generality in cell and developmental biology. Cell Syst., 15:790–807, 2024. doi: 10.1016/j.cels.2024.08.001.

[89] J. E. Fewell, A. A. Hislop, J. A. Kitterman, and P. Johnson. Effect of tracheostomy on lung development in fetal lambs. J. Appl. Physiol., 55:1103–1108, 1983. doi: 10.1152/jappl.1983.55.4.1103.

[90] Alejandro Aguilera-Castrejon, Bernardo Oldak, Tom Shani, Nadir Ghanem, Chen Itzkovich, Sharon Slomovich, Shadi Tarazi, Jonathan Bayerl, Valeriya Chugaeva, Muneef Ayyash, et al. Ex utero mouse embryogenesis from pre-gastrulation to late organogenesis. Nature, 593:119–124, 2021. doi: 10.1038/s41586-021-03416-3.

[91] T. H. Vu, Y. Alemayehu, and Z. Werb. New insights into saccular development and vascular formation in lung allografts under the renal capsule. Mech. Dev., 120:305–313, 2003. doi: 10.1016/s0925-4773(02)00451-3.

[92] Che-Hang Yu, Yiyi Yu, Liam M Adsit, Jeremy T Chang, Jad Barchini, Andrew H Moberly, Hadas Benisty, Jinkyung Kim, Brent K Young, Kathleen Heng, et al. The cousa objective: a long-working distance air objective for multiphoton imaging in vivo. Nat. Methods, 21:132–141, 2024. doi: 10.1038/s41592-023-02098-1.

[93] R. Revellin, F. Rousset, D. Baud, and J. Bonjour. Extension of Murray’s law using a non-Newtonian model of blood flow. Theor. Biol. Med. Model., 6:7, 2009. doi: 10.1186/1742-4682-6-7.

[94] P. R. Painter, P. Edén, and H.-U. Bengtsson. Pulsatile blood flow, shear force, energy dissipation and Murray’s Law. Theor. Biol. Med. Model., 3:31, 2006. doi: 10.1186/1742-4682-3-31.

[95] S. Sattari, C. A. Mariano, and M. Eskandari. Biaxial mechanical properties of the bronchial tree: Characterization of elasticity, extensibility, and energetics, including the effect of strain rate and preconditioning. Acta Biomater., 155:410–422, 2023. doi: 10.1016/j.actbio.2022.10.047.

[96] B. Suki and J. H. T. Bates. Lung tissue mechanics as an emergent phenomenon. J. Appl. Physiol., 110:1111–1118, 2011. doi: 10.1152/japplphysiol.01244.2010.

[97] J. B. West. Distribution of mechanical stress in the lung, a possible factor in localisation of pulmonary disease. Lancet, 297:839–841, 1971. doi: 10.1016/s0140-6736(71)91501-7.

[98] D. Doorly and S. Sherwin. Geometry and flow, chapter 5, pages 177–209. Springer, 2009. doi: 10.1007/978-88-470-1152-65.

[99] H. F. Gómez, N. Doumpas, and D. Iber. Time-lapse and cleared imaging of mouse embryonic lung explants to study three-dimensional cell morphology and topology dynamics. STAR Protoc., 4:102187, 2023. doi: 10.1016/j.xpro.2023.102187.

[100] M. D. Muzumdar, B. Tasic, K. Miyamichi, L. Li, and L. Luo. A global double-fluorescent Cre reporter mouse. Genesis, 45:593–605, 2007. doi: 10.1002/dvg.20335.

[101] E. A. Susaki, K. Tainaka, D. Perrin, H. Yukinaga, A. Kuno, and H. R. Ueda. Advanced CUBIC protocols for whole-brain and whole-body clearing and imaging. Nat. Protoc., 10:1709–1727, 2015. doi: 10.1038/nprot.2015.085.

[102] F. F. Voigt, D. Kirschenbaum, E. Platonova, S. Pagés, R. A. A. Campbell, et al. The mesoSPIM initiative: open-source light-sheet microscopes for imaging cleared tissue. Nat. Methods, 16:1105–1108, 2019. doi: 10.1038/s41592-019-0554-0.

[103] D. Hörl, F. Rojas Rusak, F. Preusser, P. Tillberg, N. Randel, et al. BigStitcher: reconstructing high-resolution image datasets of cleared and expanded samples. Nat. Methods, 16:870–874, 2019. doi: 10.1038/s41592-019-0501-0.

[104] J. Ahlers, D. Althviz Moré, O. Amsalem, A. Anderson, G. Bokota, et al. napari: a multi-dimensional image viewer for Python, 2023.

[105] R. Haase, K. Yamauchi, J. Müller, and I. Fernando. haesleinhuepf/apoc: 0.12.0, 2022.

[106] Comet Technologies Canada Inc. Dragonfly 2024.1, 2024. https://dragonfly.comet.tech.

[107] R. Girshick. Fast r-cnn. 1504.08083, 2015. doi: 10.48550/arXiv.1504.08083.

[108] K. Yamauchi, R. Haase, and P. Sobolewski. morphometrics/morphometrics: v0.0.9, 2023.

[109] J. A. Park, J. J. Fredberg, and J. M. Drazen. Putting the Squeeze on Airway Epithelia. Physiology (Bethesda), 30:293–303, 2015. doi: 10.1152/physiol.00004.2015.

[110] J. E. Hiorns, O. E. Jensen, and B. S. Brook. Static and dynamic stress heterogeneity in a multiscale model of the asthmatic airway wall. J. Appl. Physiol., 121:233–247, 2016. doi: 10.1152/japplphysiol.00715.2015.

[111] A. S. LaPrad, T. L. Szabo, B. Suki, and K. R. Lutchen. Tidal stretches do not modulate responsiveness of intact airways in vitro. J. Appl. Physiol., 109:295–304, 2010. doi: 10.1152/japplphysiol.00107.2010.

[112] C. M. Waters, E. Roan, and D. Navajas. Mechanobiology in lung epithelial cells: measurements, perturbations, and responses. Compr. Physiol., 2:1–29, 2012. doi: 10.1002/cphy.c100090.

[113] V. K. Sidhaye, K. S. Schweitzer, M. J. Caterina, L. Shimoda, and L. S. King. Shear stress regulates aquaporin-5 and airway epithelial barrier function. Proc. Natl. Acad. Sci. U. S. A., 105:3345–3350, 2008. doi: 10.1073/pnas.0712287105.

[114] G. Xia, M. H. Tawhai, E. A. Hoffman, and C. L. Lin. Airway wall stiffening increases peak wall shear stress: a fluid-structure interaction study in rigid and compliant airways. Ann. Biomed. Eng., 38:1836–1853, 2010. doi: 10.1007/s10439-010-9956-y.

[115] B. Sul, A. Wallqvist, M. J. Morris, J. Reifman, and V. Rakesh. A computational study of the respiratory airflow characteristics in normal and obstructed human airways. Comput. Biol. Med., 52:130–143, 2014. doi: 10.1016/j.compbiomed.2014.06.008.

